# The E3 Ubiquitin Ligase Mindbomb1 controls zebrafish Planar Cell Polarity

**DOI:** 10.1101/2021.07.05.451064

**Authors:** Vishnu Muraleedharan Saraswathy, Priyanka Sharma, Akshai Janardhana Kurup, Sophie Polès, Morgane Poulain, Maximilian Fürthauer

**Author notes:** Corresponding author: Maximilian Fürthauer. These authors contributed equally to the present study (Priyanka Sharma, Akshai Janardhana Kurup).

## Abstract

Vertebrate Delta/Notch signaling involves multiple ligands, receptors and transcription factors. Delta endocytosis – a critical event for Notch activation – is however essentially controlled by the E3 Ubiquitin ligase Mindbomb1 (Mib1). Due to its position at a molecular bottleneck of the pathway, Mib1 inactivation is often used to inhibit Notch signaling. However, recent findings indicate that the importance of Mib1 extends beyond the Notch pathway. We report an essential role of Mib1 in Planar Cell Polarity (PCP).

*mib1* null mutants or morphants display impaired gastrulation stage Convergence Extension (CE) movements. Comparison of different *mib1* mutants and functional rescue experiments indicate that Mib1 controls CE independently of Notch. In contrast, Mib1-dependent CE defects can be rescued using the PCP downstream mediator RhoA. Mib1 regulates CE through the RING Finger domains that have been implicated in substrate ubiquitination, suggesting that Mib1 may control PCP protein trafficking. Accordingly, we show that Mib1 controls the endocytosis of the PCP component Ryk and that Ryk internalization is required for CE.

Numerous morphogenetic processes involve both Notch and PCP signaling. We show that Mib1, a known Notch signaling regulator, is also an essential PCP pathway component. Care should therefore be taken when interpreting Mib1 loss of function phenotypes.

## INTRODUCTION

Endocytic membrane trafficking is essential to control the abundance, localization and activity of cellular signaling molecules. Depending on the biological context, the internalization of proteins from the cell surface allow desensitization to extracellular stimuli, formation of endosomal signaling compartments, resecretion of signaling molecules through endosomal recycling or their lysosomal degradation (Hupalowska & Miaczynska, 2012; Villaseñor et al., 2016). One key example for the importance of endosomal membrane trafficking is provided by the Delta/Notch signaling pathway, where Delta ligand endocytosis is required for Notch receptor activation (Chitnis, 2006; Fürthauer & González-Gaitán, 2009; Le Borgne, Bardin, et al., 2005; Seib & Klein, 2021).

Notch receptors are single-pass transmembrane proteins that interact with Delta/Serrate/Lag2 (DSL) family ligands (Bray, 2016; Hori et al., 2013). Productive ligand/receptor interactions trigger a series of proteolytic cleavages that release the Notch Intracellular Domain (NICD) into the cytoplasm, allowing it thereby to enter the nucleus and associate with transcriptional cofactors to activate target gene expression. Studies in *Drosophila* and vertebrates revealed that Ubiquitin-dependent endocytosis of DSL ligands in signal-sending cells is essential to promote Notch receptor activation in adjacent signal-receiving cells (Deblandre et al., 2001; Itoh et al., 2003; Lai et al., 2001; Le Borgne & Schweisguth, 2003; Pavlopoulos et al., 2001).

While the precise mechanism through which Delta promotes Notch activation is still under investigation, current models suggest that the endocytosis of DSL ligands represents a force-generating event that physically pulls on Notch receptors to promote their activation (Langridge & Struhl, 2017; Meloty-Kapella et al., 2012; Seib & Klein, 2021). Notch pathway activation is therefore critically dependent on Delta endocytosis, a process controlled by ligand poly-ubiquitination. Protein ubiquitination involves ubiquitin-activating E1 enzymes, ubiquitin-conjugating E2 enzymes and substrate-specific E3 ubiquitin ligases (Oh et al., 2018). Through their ability to recognize specific substrates, different E3 ligases control the activity of various cellular signaling pathways.

Genetic studies revealed that two different RING (Really Interesting New Gene) finger domain E3 ligases, Neuralized (Neur) and Mindbomb (Mib) control DSL ligand endocytosis in *D*rosophila (Lai et al., 2001; Le Borgne, Remaud, et al., 2005; Le Borgne & Schweisguth, 2003; Pavlopoulos et al., 2001). Vertebrate genomes harbor two *mib* and several *neur* homologues. However, mouse *neur1* or *neur2* single and double mutants present no phenotypes indicative of defective Notch signaling (Koo et al., 2007). In contrast, Notch signaling is severely impaired upon genetic inactivation or morpholino knock-down of *mib1* in mice, Xenopus and zebrafish (Itoh et al., 2003; Koo et al., 2007; Yoon et al., 2008). In addition to Mib1, its orthologue Mib2 as well as Asb11, a component of a multisubunit Cullin E3 ligase complex, have been implicated in DSL ligand endocytosis (Sartori da Silva et al., 2010; Zhang, Li, & Jiang, 2007). Nonetheless, mutational analysis in zebrafish failed to confirm a requirement for *mib2* in Notch signaling (Mikami et al., 2015) and showed that a *mib1* inactivation is sufficient to essentially abolish the expression of a transgenic Notch reporter in the central nervous system (Sharma et al., 2019). These findings identify Mib1 as the major regulator of vertebrate DSL ligand endocytosis.

In addition to Delta ligands, Mib1 is able to interact with a number of additional protein substrates (Berndt et al., 2011; Matsuda et al., 2016; Mertz et al., 2015; Tseng et al., 2014). Accordingly, functional studies have implicated Mib1 in a growing number of functions that include the regulation of epithelial morphogenesis (Dho et al., 2019; Matsuda et al., 2016) and cell migration (Mizoguchi et al., 2017), centrosome and cilia biogenesis (Douanne et al., 2019; Joachim et al., 2017; Villumsen et al., 2013; Wang et al., 2016; Čajánek et al., 2015), the control of glutamate receptor localization (Sturgeon et al., 2016) or interferon production (Li et al., 2011).

A study in human cell culture identified the Receptor-like tyrosine kinase Ryk as a target of Mib1-mediated ubiquitination (Berndt et al., 2011). Ryk is a single-pass transmembrane protein that binds Wnt ligands through its extracellular Wnt Inhibitory Factor (WIF) domain. While an intracellular pseudokinase domain appears devoid of functional enzymatic activity, Ryk has been suggested to regulate cell signaling through scaffolding functions or the γ-secretase dependent release and nuclear translocation of its intracellular domain (Roy et al., 2018). A number of studies have implicated Ryk in canonical, β-catenin dependent Wnt signaling (Green et al., 2008; Lu et al., 2004; Roy et al., 2018). In this context, Mib1 has been shown to promote the ubiquitination and internalization of Ryk, which appears to be required for Wnt3A-mediated β-catenin stabilization/activation (Berndt et al., 2011). While experiments in *C*.*elegans* provided evidence for genetic interactions between *mib1* and *ryk* (Berndt et al., 2011), the importance of Mib1/Ryk interactions for vertebrate development or physiology has not been addressed.

In addition to its role in canonical Wnt signaling, several studies have linked Ryk to the non-canonical, β-catenin independent, Wnt/Planar Cell Polarity (PCP) and Wnt/Ca^2+^ pathways (Duan et al., 2017; Kim et al., 2008; Lin et al., 2010; Macheda et al., 2012; Roy et al., 2018). *Ryk* mutant mice present a range of diagnostic PCP phenotypes, including defects in neural tube closure and the orientation of inner ear sensory hair cells (Andre et al., 2012; Macheda et al., 2012). During early fish and frog development, PCP signaling regulates Wnt-dependent Convergent Extension (CE) movements that direct embryonic axis extension (Butler & Wallingford, 2017; Davey & Moens, 2017; Gray et al., 2011; Tada & Heisenberg, 2012). Different studies have suggested that Ryk may control PCP by acting together with Wnt-binding Frizzled (Fz) receptors (Kim et al., 2008), regulating a Fz-independent parallel pathway (Lin et al., 2010) or by controlling the stability of the core PCP pathway component Vangl2 (Andre et al., 2012).

In both vertebrates and invertebrates, the core PCP machinery is defined by three transmembrane proteins Flamingo(Fmi)/CELSR, Fz and Strabismus/Vangl as well as their cytoplasmic partners Dishevelled (Dvl), Prickle (Pk) and Diego (Dgo)/ANKRD6 (Butler & Wallingford, 2017; Devenport, 2014; Harrison et al., 2020; Humphries & Mlodzik, 2018; Vladar et al., 2009). Polarity is established through formation of distinct CELSR/Vangl/Pk and CELSR/Fz/Dvl/Dgo complexes at the opposite sides of the cell. The fact that similar defects are often observed upon overexpression or inactivation of PCP pathway components suggests that the levels of individual proteins need to be tightly controlled. It is therefore no surprise that factors such as Dynamin, which governs endocytic vesicle scission, the early endosomal GTPase Rab5 and other regulators of membrane trafficking have been shown to control the distribution of the transmembrane proteins Fmi/CELSR, Fz and Vangl2 (Butler & Wallingford, 2017; Devenport et al., 2011; Mottola et al., 2010; Strutt & Strutt, 2008). In addition to regulating the trafficking of core pathway components, endocytosis ensures the PCP-dependent regulation of cellular adhesion molecules (Classen et al., 2005; Ulrich et al., 2005).

While numerous studies therefore indicate a central role of membrane trafficking in PCP, it remains to be established whether Ryk/PCP signaling is subject to endo-lysosomal control. In the present study, we show that Mib1-mediated Ryk endocytosis is required for the PCP-dependent control of CE movements during zebrafish gastrulation. The analysis of different *mib1* mutant alleles shows that the role of Mib1 in PCP is separable from its function in Notch signaling. Our work thereby identifies a novel function of this E3 ubiquitin ligase and establishes Mib1 as an essential regulator of the PCP pathway.

## RESULTS

### Mindbomb1 regulates Convergent Extension independently of Notch

Through its ability to promote Delta ligand endocytosis, the E3 Ubiquitin ligase Mindbomb1 plays an essential role in vertebrate Notch receptor activation (Guo et al., 2016). In the course of experiments that were initially designed to study the role of Notch signaling in the morphogenesis of the zebrafish nervous system (Sharma et al., 2019), we realized that embryos injected with a mib1 morpholino (mib1 morphants) present a reduced axial extension at the end of gastrulation that is indicative of defects in embryonic Convergence Extension (CE) movements (Fig.1A). Accordingly, 2 somite stage mib1 morphants present a widening of the notochord, somites and neural plate (Fig.1B). The mib1 exon/intron1 splice site morpholino used in these experiments has been previously validated in different studies (Itoh et al., 2003; Sharma et al., 2019). We further confirmed its specificity by showing that the co-injection of a WT mib1 RNA that is not targetable by the morpholino restores axis extension (Fig.1A).

**Figure 1:**
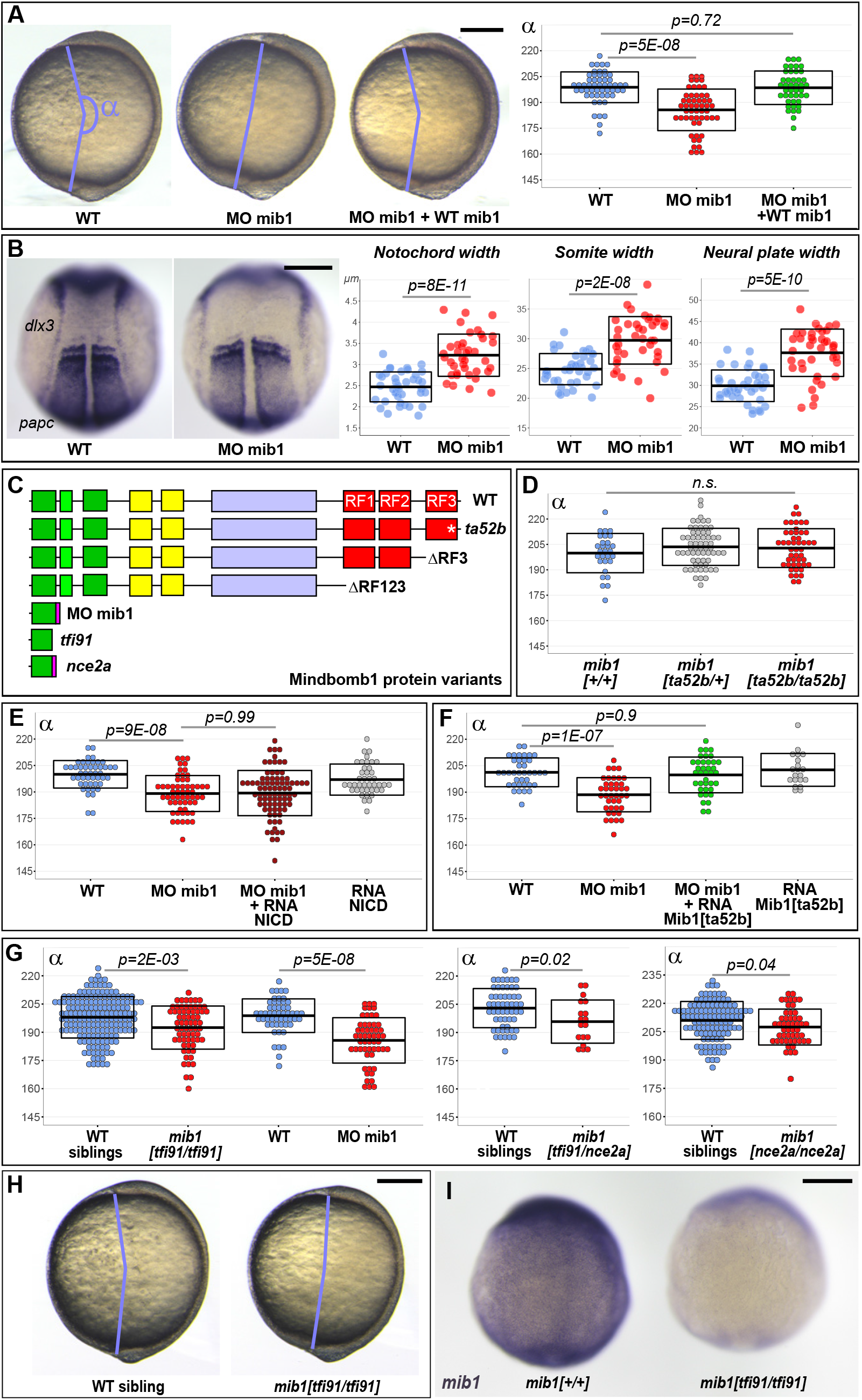
Mib1 regulates PCP-dependent convergent extension movements independently of Notch. **(A)** Axis extension was quantified at bud stage by measuring the axis extension angle α. Axis extension is reduced in mib1 morphants but restored upon coinjection of WT mib1 RNA. Lateral views of bud stage embryos, anterior up, dorsal to the right. **(B)** mib1 morphants present a widening of the notochord, somites and neural plate. Dorsal views of 2 somite stage embryos, anterior up. *dlx3 in situ* hybridization outlines the neural plate, *papc* the somites and the adaxial cells lining the notochord. Widths indicated in microns. **(C)** Mib1 protein variants used in the study. **(D)** The *mib1*^*ta52b*^ mutation has no effect on axis extension. **(E)** Constitutively activated Notch (NICD) fails to restore mib1 morphant axis extension. **(F)** mib1^ta52b^ RNA injection restores mib1 morphant axis extension. **(G,H)** Axis extension is impaired in *mib1*^*tfi91*^ or *mib1*^*nce2a*^ null mutants. On the left panel the mib1 morphant data from Fig. 1A are included for comparison. **(I)** *In situ* hybridization reveals reduced *mib1* transcript levels in n=27 *mib1*^*tfi91*^ mutant embryos. Dorsal views of bud stage embryos, anterior up. To warrant identical acquisition conditions, two embryos were photographed on a single picture. Scalebars: 200 µm. Boxes in (A,B, D-G) represent mean values +/- SD. See supplementary material for complete statistical information.

The *mib1*^*ta52b*^ mutation in the C-terminal Mib1 RING finger domain (RF3, Fig.1C) disrupts the ability of the protein to promote Delta ubiquitination and thereby disrupts Notch signaling (Itoh et al., 2003; Sharma et al., 2019; Zhang, Li, & Jiang, 2007). Axial extension occurs however normally in *mib1*^*ta52b*^ mutants (Fig.1D, Suppl.Fig.1A), raising the question whether Mib1 exerts a Notch-independent function in CE. In accordance with this hypothesis, a constitutively activated form of Notch (NICD) that is able to restore Notch-dependent defects in the nervous system (Sharma et al., 2019) fails to rescue mib1 morphant axis extension (Fig.1E, Suppl.Fig.2A).

Sequencing of the *mib1* cDNA in mib1 morphants revealed a retention of intron1 that causes the appearance of a premature Stop codon (Suppl.Fig.2B). As a consequence, the Mib1 morphant protein comprises only the first 76 amino acids of WT Mib1 (Fig.1C). We hypothesized that this early termination of the open reading frame could disrupt functions that are not affected by the *mib1*^*ta52b*^ point mutation. Accordingly, mib1 WT and *mib1*^*ta52b*^ mutant RNAs are equally capable of rescuing mib1 morphant CE phenotypes (Fig.1A,F, Suppl.Fig.2C).

As *mib1*^*ta52b*^ point mutants show normal CE, we further studied axis extension in *mib1* null mutants. The previously reported *mib1*^*tfi91*^ allele causes a truncation of the Mib1 open reading frame after 59 amino acids and therefore likely represents a molecular null (Fig. 1C) (Itoh et al., 2003). In contrast to *mib1*^*ta52b*^ mutants, *mib1*^*tfi91*^ homozygous animals present CE defects that are statistically significant, although weaker than in mib1 morphants (Fig.1G,H, Cohen’s d effect size = 0.49 for *mib1*^*tfi91*^, 1.23 for MO mib1). Similar phenotypes are observed for a new potential null allele generated in the present study (*mib1*^*nce2a*^, Fig.1C,G, Suppl.Fig.1C,E) or in *mib1*^*tfi91/nce2a*^ trans-heterozygotes (Fig.1G). The *mib1*^*tfi91*^ and *mib1*^*nce2a*^ mutations introduce stop codons shortly after the beginning of the *mib1* open reading frame ((Itoh et al., 2003) and Suppl.Fig.1E), a mutation pattern that can cause nonsense mediated decay of mutant mRNAs and could thereby trigger partial transcriptional compensation (El-Brolosy et al., 2019). Accordingly, *mib1* transcript levels appear reduced in *mib1*^*tfi91*^ and *mib1*^*nce2a*^ but not in *mib1*^*ta52b*^ mutants (Fig.1I, Suppl.Fig.1B,D).

Zebrafish Mib1 interacts with Epb41l5 to regulate neuronal differentiation (Matsuda et al., 2016) and with Catenin delta1 to control cell migration (Mizoguchi et al., 2017). Both of these activities are disrupted in *mib1*^*ta52b*^ mutants (Matsuda et al., 2016; Mizoguchi et al., 2017). Our observation that mib1 morphants (Fig.1A) or *mib1* null mutants (Fig.1G) but not *mib1*^*ta52b*^ mutants (Fig.1D) present defects in gastrulation stage axial extension identify thereby a novel role of Mib1 in the regulation of embryonic CE.

### Mindbomb1 RING finger domains are required for PCP

Vertebrate CE requires non-canonical Wnt/PCP signaling (Butler & Wallingford, 2017; Davey & Moens, 2017; Gray et al., 2011; Tada & Heisenberg, 2012). To test whether Mib1 loss-of-function impairs PCP pathway activity, we overexpressed the PCP downstream effector RhoA in mib1 morphants. RhoA fully restores axis extension (Fig.2A), suggesting thereby that Mib1 is required for the PCP-dependent control of embryonic CE movements.

**Figure 2:**
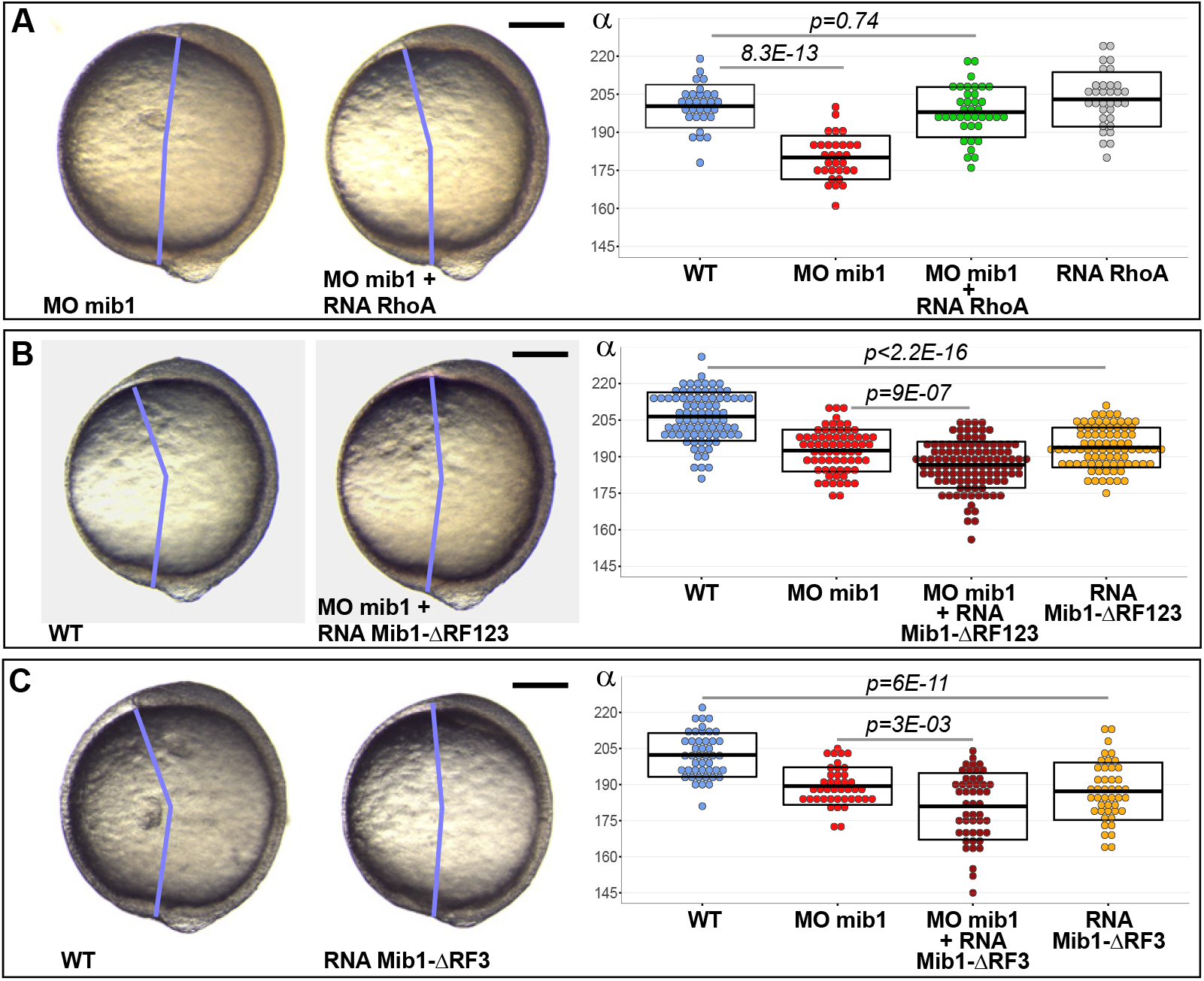
Mib1 controls PCP through its RING finger domains. **(A)** RhoA overexpression rescues mib1 morphant axis extension. **(B,C)** Mib1 proteins lacking all (Mib1ΔRF123, B) or only the last (Mib1ΔRF3, C) RING finger impair axis extension in mib1 morphant or WT embryos. Lateral views of bud stage embryos, anterior up, dorsal to the right. Scalebars: 200 µm. Boxes represent mean values +/- SD. See supplementary material for complete statistical information.

All Mib1 functions known to date require its E3 ubiquitin ligase activity that is dependent of the presence of C-terminal RING finger domains (Guo et al., 2016). In contrast to the rescuing activity of WT Mib1 (Fig.1A), a truncated Mib1 variant that lacks all three RING Finger domains (Mib1^ΔRF123^, Fig. 1C) enhances the defects of mib1 morphants as well as impairing axis extension in WT animals (Fig.2B). As RING finger deficient Mib1 variants can act as substrate-sequestering dominant negatives (Guo et al., 2016), the enhanced CE defects of Mib1^ΔRF123^-injected mib1 morphants are likely due to its capacity to interfere with maternally provided Mib1 that is not targeted by the mib1 splice morpholino. A Mib1 variant lacking only the last RING finger (Mib1ΔRF3, Fig.1C) produced similar results (Fig.2C).

Our results suggest that the substrate-ubiquitinating Mib1 RING-finger domains are required for PCP. As Ubiquitin-dependent membrane trafficking is important for PCP (Butler & Wallingford, 2017; Devenport, 2014; Feng et al., 2021), we set out to determine whether Mib1 controls CE by regulating the trafficking of a PCP pathway component.

### Convergent Extension requires Mindbomb1-dependent Ryk internalization

Mammalian Mib1 has been shown to control the ubiquitin-dependent endocytic internalization of the the Wnt co-receptor Receptor like tyrosine kinase Ryk (Berndt et al., 2011). Interestingly, studies in mice, frogs and zebrafish have implicated Ryk in non-canonical Wnt/PCP signaling (Kim et al., 2008; Lin et al., 2010; Macheda et al., 2012). To determine whether Mib1 regulates CE by controlling Ryk internalization, we started by analyzing the effect of Mib1 gain or loss of function on Ryk localization. Ryk localizes to the cell surface as well as intracellular compartments (Fig.3A), 70.7% of which are positive for the early endosomal marker Rab5 (n=75 cells from 8 embryos, Suppl.Fig.3A) (Berndt et al., 2011; Kim et al., 2008; Lin et al., 2010). In accordance with a role of Mib1 in promoting Ryk endocytosis, Mib1 overexpression depleted Ryk from the cell cortex and triggered its accumulation in intracellular compartments (Fig. 3B) without affecting the localization of the general plasma membrane marker GAP43-RFP (Suppl.Fig.3B,C).

**Figure 3:**
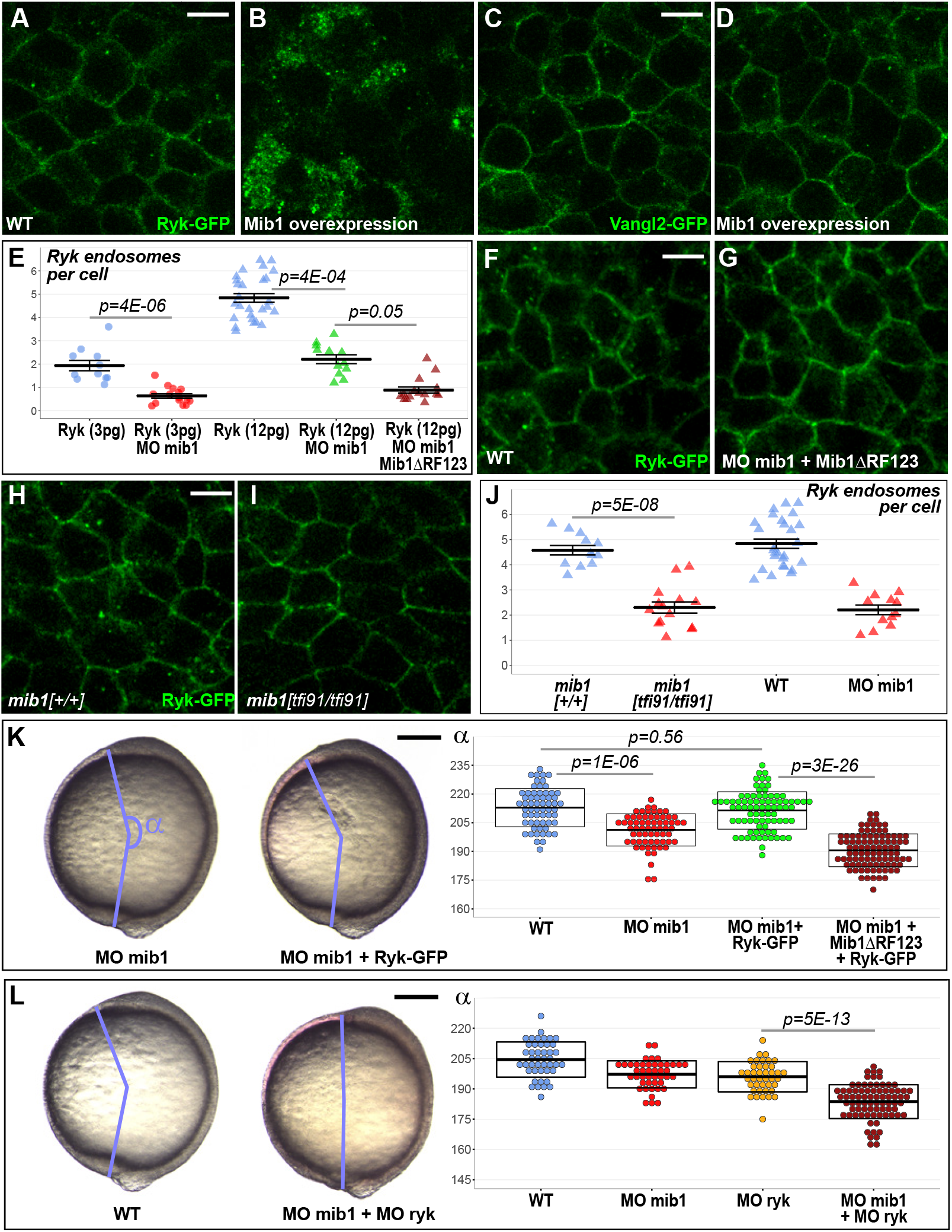
Mib1-mediated Ryk endocytosis controls Convergent Extension movements. **(A-D)** WT mib1 RNA injection triggers Ryk internalization in 20/21 embryos (B) but has no effect on Vangl2 localization (D, n=23). **(E-G)** Mib1 morpholino injection reduces the number of Ryk endosomes that are present upon injection of Ryk-GFP RNA. Increasing the dose of Ryk-GFP RNA restores endosome number in mib1 morphants but not in embryos coinjected with Mib1ΔRF123. **(H-J)** The number of Ryk endosomes that are present upon injection of Ryk-GFP RNA (12 pg) is reduced in *mib1* null mutants. mib1 morphant data from panel E are shown again for comparison. **(K)** Ryk-GFP RNA (12 pg) rescues axis extension in mib1 morphants but not in embryos coinjected with Mib1ΔRF123. **(L)** Ryk morpholino injection aggravates mib1 morphant axis extension phenotypes. (A-D,F,G,H,I) dorsal views of 90% epiboly stage embryos, anterior up, scalebars 10 µm. (K,L) Lateral views of bud stage embryos, anterior up, scalebars 200 µm. In (E,J) each data point represents the mean number of endosomes for 20 cells from a single embryo. For comparison J again includes the mib1 morphant E. Bars represent mean values +/- SEM. In (K,L) boxes represent mean values +/- SD. See supplementary material for complete statistical information.

To determine whether Mib1 specifically affects Ryk or acts as a general regulator of PCP protein trafficking, we tested the effect of Mib1 overexpression on the localization of different transmembrane proteins. Ryk has been shown to interact with the core PCP component Vangl2 whose endocytic trafficking is crucial for PCP (Andre et al., 2012). In contrast to Ryk, Mib1 overexpression has no obvious effect on Vangl2 localization (Fig. 3C,D). Similarly, we detected no impact of Mib1 overexpression on the localization of Fz2 and Fz7, two Wnt receptors that have been in implicated in PCP signaling (Kim et al., 2008; Lin et al., 2010; Čapek et al., 2019) (Suppl.Fig.4A-D). Ryk belongs to a family of Wnt-binding Receptor Tyrosine Kinases that have the particularity of harboring a most likely inactive pseudokinase domain (Roy et al., 2018). Other members of this protein family include the Receptor tyrosine kinase-like Orphan Receptors (ROR), among which zebrafish ROR2 has been implicated in CE (Bai et al., 2014; Mattes et al., 2018). Our experiments failed to identify an effect of Mib1 overexpression on the localization of zebrafish ROR2 or its orthologue ROR1 (Fig.Suppl.4E-H). Taken together, our observations suggest that Mib1 regulates CE movements through the specific control of Ryk localization.

To determine if Mib1 function is required for Ryk endocytosis, we quantified the number of RYK-positive endosomes in mib1 morphants and *mib1*^*tfi91*^ null mutants. In these experiments, RNAs encoding RYK-GFP and Histone2B-mRFP were co-injected into mib1 morphant or *mib1*^*tfi91*^ mutant embryos. The Histone2B-mRFP signal was used as an injection tracer to ascertain that the number of RYK-GFP positive endosomes was scored in embryos that had received comparable amounts of injected material.

These experiments revealed a reduction in the number of Ryk-GFP positive endosomes in mib1 morphants (Fig.3E). Our observation that the CE defects of mib1 morphants can be further enhanced by the co-injection of antimorphic Mib1^ΔRF123^ or Mib1^ΔRF3^ constructs (Fig.2B,C) suggests that Mib1 activity is only partially reduced in mib1 morphants. Accordingly, increasing the amount of injected Ryk-GFP RNA from 3 to 12 pg restores the number of Ryk positive endosomes in mib1 morphants (Fig.3E). Even a high dose of Ryk-GFP fails however to restore endosome number in embryos co-injected with mib1 morpholino and dominant-negative Mib1^ΔRF123^ (Fig. 3E,F,G, Suppl.Fig5A,B), suggesting that Ryk can no more be internalized once Mib1 function is severely compromised.

Similar to mib1 morphants, *mib1*^*tfi91*^ null mutants present a reduction in the number of RYK-GFP positive endosomes (Fig.3H,I,J, Suppl.Fig.5C,D). While stronger CE defects are observed in mib1 morphants compared to *mib1* mutants (Fig.1G), morphant and mutant embryos present a comparable reduction of the number of RYK-GFP positive endosomes (Fig.3J, Cohen’s d effect size = 3.02 for *mib1*^*tfi91*^, 2.99 for mib1 morphants). These observations suggest the existence of a compensatory mechanism that allows to partially correct the PCP signaling defects that arise from a defect in Ryk endocytosis.

Why does impaired Ryk endocytosis cause CE defects? The phenotypes of Mib1-depleted embryos could be either due to the loss of Ryk-positive endosomal compartments, or to the accumulation of non-internalized Ryk at the cell surface. In *C*.*elegans*, Mib1 controls Ryk cell surface levels by promoting Ryk internalization and degradation (Berndt et al., 2011). Similarly, zebrafish embryos injected with a high level of Mib1 RNA present an overall loss of Ryk-GFP signal (Suppl.Fig.3D-F). If the CE phenotypes of mib1 morphants are indeed due to increased Ryk cell surface levels, then Ryk-GFP overexpression should further enhance the defects of Mib1-depleted animals. The opposite prediction should however apply if the CE defects of mib1 morphants are due to the loss of Ryk-positive endosomal compartments. In this case Ryk-GFP overexpression, which allows to restore the number of Ryk endosomes in mib1 morphants (Fig.3E), should also rescue the CE phenotypes of Mib1-depleted animals. In accordance with this later hypothesis, Ryk-GFP overexpression restores axis extension (Fig.3K). In contrast, Vangl2 overexpression did not display rescuing activity (Suppl.Fig.6).

If mib1 morphant CE defects are due to a loss of Ryk-positive endosomes, ryk knock-down is expected to further increase the severity of the observed phenotypes. To test this hypothesis, morpholinos directed against mib1 or ryk were injected separately or in combination. The injection of morpholino-insensitive mib1 or ryk RNAs allows to rescue the CE defects that are generated by their respective morpholinos, validating thereby the specificity of the reagents (Fig.1A, Fig.4D, Suppl.Fig.7A). Co-injection of mib1 and ryk morpholinos causes CE defects that are significantly enhanced compared to single morphants (Fig.3L), adding further support to our hypothesis that the CE defect of Mib1-depleted embryos are due to Ryk loss of function.

**Figure 4:**
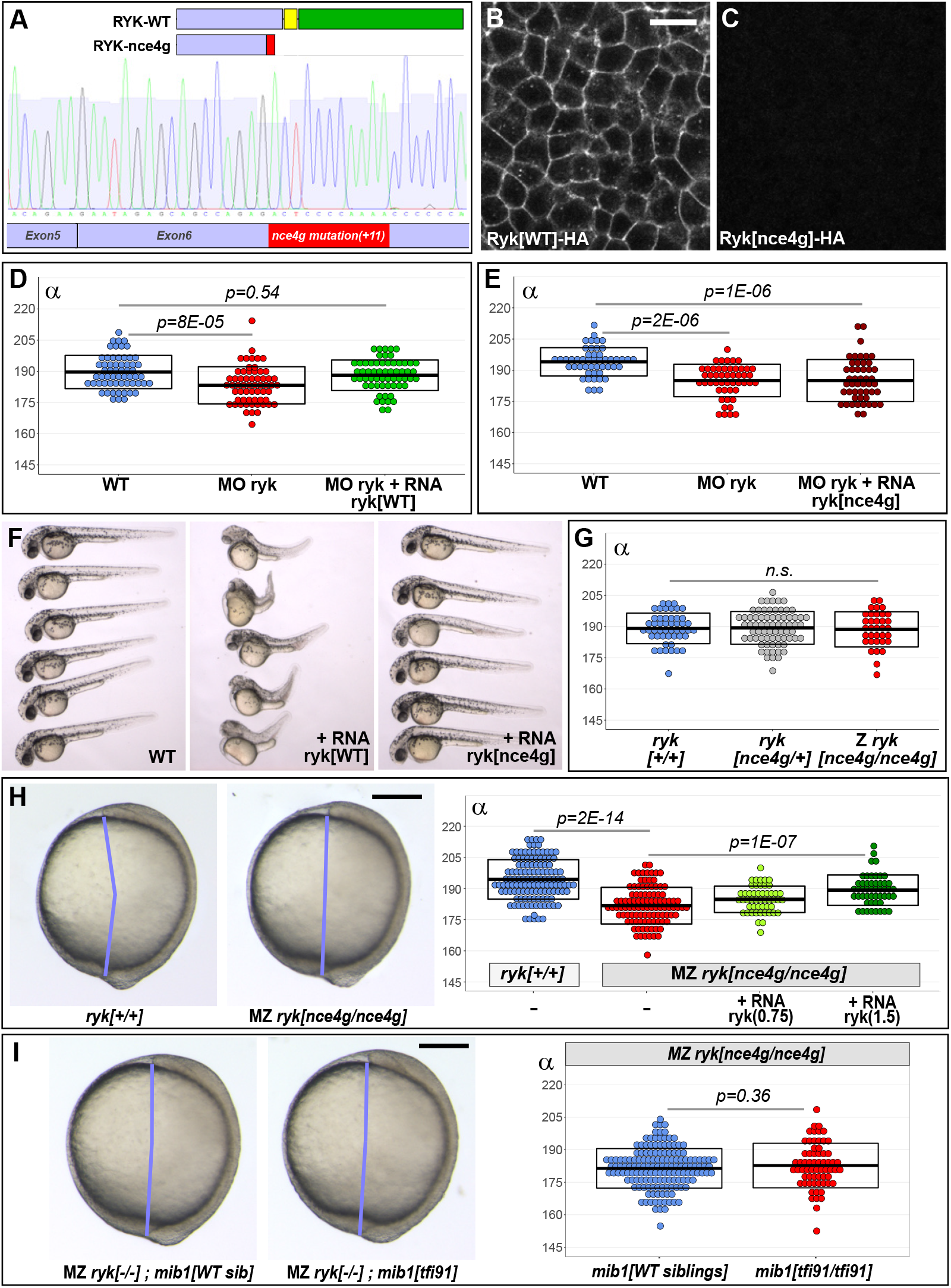
*mindbomb1* loss of function has no effect on convergent extension in maternal zygotic *ryk* mutants. **(A)** *ryk*^*nce4g*^ mutants present an 11 base pair insertion in exon 6. The RYK-nce4g mutant protein comprises only a part of the extracellular (blue) and lacks the entire transmembrane (yellow) and intracellular (green) domains. **(B,C)** Accordingly, a C-terminal HA tag that allows to localize WT Ryk (B, n=12) becomes undetectable upon introduction of the *ryk*^*nce4g*^ mutation (C, n=14). Dorsal views of 90% epiboly stage embryos, anterior up. Scalebar 20 µm. **(D,E)** The Convergent Extension (CE) phenotypes of ryk morphant animals can be rescued using 1.5 pg WT ryk (D) but not *ryk*^*nce4g*^ mutant (E) RNA. **(F)** Overexpressing high levels (25 pg) WT ryk RNA causes severe embryonic malformations while no effect is observed using *ryk*^*nce4g*^ mutant RNA. 32 hpf embryos, anterior to the left, dorsal up (n=24 embryos/condition). **(G)** Zygotic *(Z) ryk* loss of function does not impair CE. **(H)** Maternal Zygotic (MZ) *ryk* mutants present defects in CE. ryk WT RNA injection allows a significant rescue of the observed phenotypes. **(I)** Similar CE defects are observed in MZ *ryk* single mutants and MZ *ryk* ; *mib1* double mutants. (H,I) Lateral views of bud stage embryos, anterior up, dorsal to the right. Scalebars 200 µm. In (D,E,G,H,I) boxes represent mean values +/- SD. See supplementary material for complete statistical information.

### *ryk* mutants are insensitive to *mindbomb1* loss of function

The abovementioned experiment shows that inhibiting Mib1 function enhances the CE defects of animals that present a partial loss of Ryk activity due to morpholino knock-down (Fig.3L). If Mib1 regulates CE by controlling Ryk endocytosis, Mib1 loss of function should however have no more enhancing effect in animals that are not only partially, but entirely devoid of Ryk activity. To test this hypothesis, we used Crispr/Cas9 mutagenesis to generate a stable *ryk* mutant line.

The use of a gRNA directed against exon 6 of zebrafish *ryk* led to the generation of *ryk*^*nce4g*^ mutants that present an 11 base pair insertion at the target locus. The presence of the mutation in *ryk*^*nce4g*^ transcripts was confirmed through sequencing of the complete *ryk* mutant cDNA (Fig.4A). The *ryk*^*nce4g*^ mutation introduces a premature stop codon that truncates the extracellular domain and deletes the transmembrane and cytoplasmic parts of the protein (Fig.4A). Accordingly, the *nce4g* mutation abolishes the detection of an HA-tag located at C-terminus of the wild-type protein (Fig.4B,C). In contrast to wild-type RNA, *ryk*^*nce4g*^ RNA fails to rescue ryk morphant CE defects (Fig.4D,E, Suppl.Fig.7A,B). While high levels of WT ryk RNA induce severe embryonic malformations, no overexpression phenotypes were observed using *ryk*^*nce4*g^ mutant RNA (Fig.4F).

Despite this evidence for functional *ryk* inactivation, our analysis failed to reveal CE defects in zygotic *ryk*^*nce4g*^ mutants (Fig.4G, Suppl.Fig.7C). Frame-shift mutations can induce nonsense mediated degradation of mutant transcripts and thereby trigger a process of transcriptional adaptation that could compensate for the loss of function of the mutated protein (El-Brolosy et al., 2019). Accordingly, *ryk* transcript levels are reduced in *ryk*^*nce4g*^ mutants (Suppl.Fig.7D).

Alternatively, the lack of CE phenotypes in zygotic *ryk*^*nce4g*^ mutants could be due to the persistence of maternally deposited *ryk* RNA or protein. *ryk*^*nce4g*^ mutants are viable and fertile and can thereby be used to generate embryos that are devoid of both maternal and zygotic Ryk function. In contrast to zygotic mutants, Maternal Zygotic (MZ) *ryk*^*nce4g*^ mutants present highly significant CE defects compared to wild-type control animals from the same genetic background (Fig.4H, see methods for details). WT ryk RNA injection induces a significant rescue of CE, confirming thereby the specificity of the observed phenotypes (Fig.4H, Suppl.Fig.7E). In further accordance with a loss of ryk activity in these animals, *ryk*^*nce4g*^ mutants are insensitive to the injection of a translation-blocking ryk morpholino that can target both maternal and zygotic RNAs (Suppl.Fig.S7F).

To determine the effect of *mib1* loss of function in embryos that are totally devoid of Ryk activity, we introduced the *mib1*^*tfi91*^ mutation in the MZ *ryk*^*nce4g*^ mutant genetic background. MZ *ryk*^*nce4g*^;*mib1*^*tfi91*^ double mutants display no significant difference in CE compared to MZ *ryk*^*nce4g*^ single mutants (Fig.4I). This observation is in agreement with our model that the genetic inactivation of *mib1* results in a specific impairment of Ryk activity. Accordingly, Mib1 loss of function has no further consequences on CE in animals that are already devoid of Ryk.

## DISCUSSION

Vertebrate Notch signaling involves multiple ligands, receptors and downstream transcription factors, but the internalization of Delta ligands that triggers Notch receptor activation is regulated essentially by Mib1 (Koo et al., 2007; Mikami et al., 2015). For this reason, numerous studies have turned to the functional inactivation of *mib1* to inhibit Notch signaling in different biological contexts. A number of recent studies have however shown that the role of Mib1 extends beyond the Notch pathway (Li et al., 2011; Mizoguchi et al., 2017; Sturgeon et al., 2016; Villumsen et al., 2013; Čajánek et al., 2015). In the present study, we have identified a novel Notch-independent function of Mib1 in the regulation of PCP-dependent CE movements during zebrafish gastrulation.

We show that two potential *mib1* null alleles, *mib1*^*tfi91*^ and the newly generated *mib1*^*nce2a*^ (which retain only the first 59 or 57 amino acids of the 1130 residue wild type protein), cause defects in gastrulation stage axis extension movements (Fig.1G). Similar defects are observed in mib1 morphants (Fig.1A) but not in *mib1*^*ta52b*^ mutants that present a missense mutation in the C-terminal RF domain (Fig.1D). In the context of Delta/Notch signaling, the Mib1^ta52b^ mutant protein exerts a dominant-negative activity and thereby causes Notch loss of function phenotypes that are even more severe than the ones observed for *mib1*^*tfi91*^ null mutants (Zhang, Li, Lim, et al., 2007). The fact that *mib1*^*ta52b*^ mutants present no defects in embryonic CE suggests therefore that the role of Mib1 in CE is distinct from its function in Notch signaling. This hypothesis is further substantiated by the finding that the CE phenotypes of mib1-depleted animals can be rescued using the PCP pathway component RhoA (Fig.2A), but not with constitutively active Notch (Fig.1E).

All known Mib1 functions require the C-terminal RF domains that are key for the protein’s E3 ligase activity. In accordance with a similar mode of action, a Mib1 variant that lacks all three RF domains is unable to support proper CE (Fig.2B). While axis extension occurs normally in *mib1*^*ta52b*^ RF3 point mutants, a Mib1 RF3 deletion variant is unable to support CE (Fig.2C). Previous studies revealed that the N-terminal part of Mib1 interacts with specific protein substrates (Berndt et al., 2011; Guo et al., 2016). Our observations show that the function of Mib1 in specific signaling pathways can also be altered through alterations in RF3. As the RF domains interact with ubiquitin-conjugating E2 enzymes (Guo et al., 2016), it is tempting to speculate that Mib1 might regulate Notch and PCP signaling using different E2 enzymes. Mib1 promotes Ryk endocytosis in mammalian cell culture (Berndt et al., 2011), but the functional relevance of this interaction for vertebrate development has not been addressed. Studies in Xenopus and zebrafish identified Ryk functions in PCP-dependent morphogenetic processes (Kim et al., 2008; Lin et al., 2010; Macheda et al., 2012). Our experiments in *mib1* null mutants and mib1 morphants identify Mib1 as an essential regulator of Ryk endocytosis and Ryk-dependent CE (Fig.3J,K).

Both *mib1* mutants and mib1 morphants present a partial impairment of Ryk endocytosis (Fig.3J). Due to this situation, we were able to restore the number of Ryk-positive endosomes by Ryk overexpression (Fig.3E). The observation that Ryk overexpression not only rescues the number of Ryk-positive endosomes but also the CE movements of mib1-depleted animals (Fig.3K) suggests that Ryk-positive endosomal compartments are required for PCP signaling.

The partial loss of Ryk-positive endosomes that is observed in *mib1* mutants is most likely due to the perdurance of maternally deposited products. Accordingly, Ryk endocytosis is nearly completely abolished in embryos that have been co-injected with mib1 splice morpholino (that blocks zygotic mib1 production) and dominant-negative Mib1^ΔRF123^ (that blocks maternal protein). The observation that under these conditions neither the number of Ryk-positive endosomes (Fig.3E), nor embryonic CE (Fig.3K) can be restored by Ryk overexpression suggests that Ryk endocytosis is essentially dependent on Mib1.

As a growing number of Mib1 substrates are being identified (Matsuda et al., 2016; Mertz et al., 2015; Mizoguchi et al., 2017; Tseng et al., 2014), the question arises whether Mib1 could regulate the ubiquitin-dependent trafficking of additional PCP pathway components. Our observations do not support this hypothesis: First, Mib1 overexpression specifically promotes Ryk internalization, while having no discernable effect on the localization of other PCP-related transmembrane proteins (Fig.3A-D, Suppl.Fig.4). Second, our analysis of *mib1;ryk* double mutants shows that the CE defects that are already observed in animals that are entirely devoid of Ryk are not further enhanced by the loss of Mib1 (Fig.4I).

Despite the fact that *mib1* null mutants and mib1 morphants present a similar reduction in the number of Ryk-positive endosomes (Fig.3J), CE defects are more severe in morphants than mutants (Fig.1G). The *mib1*^*tfi91*^ and *mib1*^*nce2a*^ alleles used in our study introduce early stop codons in the *mib1* open reading frame, a mutation pattern that has been reported to induce degradation of mutant transcripts and upregulation of compensatory genes (El-Brolosy et al., 2019). In accordance with such a scenario, *mib1* transcript levels are reduced in these *mib1* mutants (Fig.1I, Suppl.Fig.1D). While compensation could potentially occur through a mechanism that promotes Mib1-independent Ryk endocytosis, our data do not support such a hypothesis (Fig.3J). Instead, our observations indicate a mechanism of PCP pathway resilience that allows to partially correct the defects that arise from a failure in Ryk endocytosis.

Taken together, our findings identify the E3 ubiquitin ligase Mib1 as an essential novel PCP regulator. Mib1 regulates CE movements through the control of Ryk endocytosis, independent of its role in Delta/Notch signaling. As processes such as the morphogenesis of the vertebrate neural tube involve both Notch and PCP signaling, our data suggest that great care should be taken for the interpretation of Mib1 loss of function phenotypes.

## MATERIALS AND METHODS

### Crispr/Cas mutagenesis

#### Generation of a *mib1*^*nce2a*^ mutants

Crispr/Cas9 mutagenesis of zebrafish *mib1* was performed using the reverse strand exon 1 target site 5’-GGAGCAGCGGTAATTGGCGG**CGG**-3’ (bold lettering indicates the PAM motif). gRNA design and *in vitro* transcription were performed according to reported protocols (Hruscha et al., 2013; Jao et al., 2013). A pre-assembled complex of purified Cas9 protein (NEB) and gRNA was injected and the efficiency of Crispr/Cas9-induced mutagenesis in the F0 generation monitored at 24 hpf using a T7 endonuclease assay (Jao et al., 2013) on a PCR amplicon comprising the Crispr target region (Forward primer: 5’-TGACTGGAAGTGGGGGAAGC-3’, Reverse primer: 5’-TGCAGTATTAGAAACGCGTG-3’). Direct sequencing of the same PCR amplicon was used to identify induced mutations in the F1 generation. This procedure led to the identification of the *mib1*^*nce2a*^ mutant allele which introduces a frame shift after amino acid 57 and induces the appearance of a premature stop codon after residue 69.

#### Generation of a *ryk*^*nce4g*^ mutants

Crispr/Cas9 mutagenesis of zebrafish *ryk* was performed using the reverse strand exon 5 target site 5’-GGCAGAGTTTTGGGGGGCTC**TGG**-3’ using the strategy mentioned above. The T7 endonuclease assay in the F0 generation and mutation identification in F1 were performed using the same PCR amplicon (Forward primer: 5’-GTGATGTTAGACTTGCATAC-3’, Reverse primer: 5’-GAAGGTTTACAAGGGCAGAATG-3’). The *ryk*^*nce4g*^ mutation introduces an 11 bp insertion that causes a frame shift after amino acid 196 and induces the appearance of a premature stop codon after residue 214.

### Fish strains and molecular genotyping

Unless otherwise specified, experiments were performed in embryos derived from an ABTÜ hybrid wild-type strain. Mutant strains included *mib1*^*ta52b*^ (Itoh et al., 2003), *mib1*^*tfi91*^ (Itoh et al., 2003), *mib1*^*nce2a*^ (this study) and *ryk*^*nce4g*^ (this study).

Depending on the experiment, *mib1* homozygous mutants were identified using molecular genotyping (see below) or through the identification of the characteristic white tail phenotype that can easily be identified by 36 hpf.

For the genotyping of *mib1*^*ta52b*^ mutants a 4-primer-PCR was used to identify WT and mutant alleles in a single PCR. The primers used were: 5’-ACAGTAACTAAGGAGGGC-3’ (generic forward primer), 5’-AGATCGGGCACTCGCTCA-3’ (specific WT reverse primer), 5’-TCAGCTGTGTGGAGACCGCAG-3’ (specific forward primer for the *mib1*^*ta52b*^ allele), and 5’-CTTCACCATGCTCTACAC-3’ (generic reverse primer). WT and *mib1*^*ta52b*^ mutant alleles respectively yield 303 bp and 402 bp amplification fragments. As some zebrafish strains present polymorphic *mib1* WT alleles, it is important to validate the applicability of this protocol before use in a given genetic background.

For the genotyping of *mib1*^*tfi91*^ mutants two separate allele-specific PCRs were used to identify WT and mutant alleles. The primers used were: 5’-TAACGGCACCGCCGCCAATTAC-3’ and 5’-GCGACCCCAGATTAATAAAGGG-3’ (WT allele), 5’-ATGACCACCGGCAGGAATAACC-3’ and 5’-ACATCATAAGCCCCGGAGCAGCGC-3’ (mutant allele).

For the genotyping of *mib1*^*nce2a*^ mutants two separate allele-specific PCRs were used to identify WT and mutant alleles. The primers used were: 5’-GCAGGAATAACCGAGTGATG-3’ and 5’-AGCAGCGGTAATTGGCGG-3’ (WT allele), 5’-GCAGGAATAACCGAGTGATG-3’ and 5’-GAGCAGCGGTAATTGAATA-3’ (mutant allele).

For the genotyping of *ryk*^*nce4g*^ mutants a single PCR reaction was used to amplify the mutation-carrying region (Forward primer 5’-GTGATGTTAGACTTGCATAC-3’, Reverse primer 5’-GAAGGTTTACAAGGGCAGAATG-3’). Due to the presence of an 11 bp insertion, mutant and WT alleles can be distinguished on a 2.5% agarose gel.

To avoid issues related to variations among genetic backgrounds, the different adult fish used in the course of our analysis of *ryk*^*nce4g*^ single and *ryk*^*nce4g*^ ; *mib1*^*tfi91*^ double mutants (Fig.4G,H,I, Suppl.Fig.7C,E,F) were all derived from a single incross of *ryk*^*nce4g/+*^ ; *mib1*^*tfi91/+*^ parents.

For DNA extraction embryos were lysed 20 min at 95°C in 28.5 µl 50 mM NaOH, and then neutralized by adding 1.5 µl Tris-HCl pH 7.5. PCR amplifications were carried out using GoTaq G2 polymerase (Promega) at 1.5 mM MgCl2 using the following cycling parameters: 2 min 95°C - 10 cycles [30 sec 95°C – 30 sec 65 to 55°C – 60 sec 72°C] – 25 cycles [30 sec 95°C – 30 sec 55°C – 60 sec 72°C] – 5 min 72°C.

### mRNA and morpholino injections

Microinjections into dechorionated embryos were carried out using a pressure microinjector (Eppendorf FemtoJet). Capped mRNAs were synthesized using the SP6 mMessage mMachine kit (Ambion). RNA and morpholinos were injected together with 0.2% Phenol Red. Morpholinos were injected at 500 (mib1 5’-GCAGCCTCACCTGTAGGCGCACTGT-3’, (Itoh et al., 2003)) or 62.5 µM (ryk 5’-GGCAGAAACATCACAGCCCACCGTC-3’).

RNA microinjection was performed using the following constructs and quantities: Mib1^ta52b^-pCS2+ (125 pg) and Mib1^ΔRF123^ (125 pg) (Zhang, Li, & Jiang, 2007). Mib1-pCS2+ (12.5 pg unless otherwise indicated) and Mib^ΔRF3^-pCS2+ (125 pg) (this study). Ryk-GFP-pCS2+ (25 pg) and Flag-Myc-Ryk-pCS2+ (50 pg) (Lin et al., 2010). Ryk-pCS2+ (0.75-25 pg), Ryk^nce4g^-pCS2+ (1.5-25 pg), Ryk-HA-pCS2+ (25 pg), Ryk^nce4g^-HA-pCS2+ (25 pg), Ryk-GFP-pCS2+ (3-12 pg) (this study, all constructs have been engineered to abolish ryk morpholino binding without altering the Ryk protein sequence). GFP-Vangl2-pCS2+ (Mahuzier et al., 2012). Fz2-mCherry-pCS2+ (50 pg) (Lin et al., 2010). Fz7-YFP-pCS2+ (25 pg) (Witzel et al., 2006). ROR1-GFP-pCS2+ (25 pg, this study). ROR2-mCherry-pCS2+ (25 pg) (Mattes et al., 2018). GFP-Rab5c-pCS2+ (50 pg, this study). RhoA-pCS2+ (25 pg) (Castanon et al., 2013). NICD-pCS2+ (37.5 pg) (Takke & Campos-Ortega, 1999). GAP43-RFP-pCS2+ (25 pg). Histone2B-mRFP-pCS2+ (12.5 pg) (Gong et al., 2004). Histone2B-GFP (6 pg) and Histon2B-tagBFP (1.5 pg) (this study).

### RNA *in situ* hybridization

Whole mount RNA *in situ* hybridizations were performed as previously described (Thisse & Thisse, 2008). The *dlx3* probe has been previously described (Kilian et al., 2003). *ryk* antisense RNA was transcribed from ryk-pBSK (this study). For *papc* and *mib1 in situ* probes were transcribed from PCR products that contained a T7 promoter sequence at their 3’end. The papc region amplified from genomic DNA extended from 5’-TCCTTCTGCAGCTCGTCCGACTGGAAG-3’ (forward strand) to 5’-GGTAAACCACCCACAGTTGAC-3’ (reverse). The mib1 probe region amplified from Mib1-pCS2+ extended from 5’-CCCGAGTGCCATGCGTGTGCTGC-3’ (forward) to 5’-CGCCGAATCCTGCTTTAC-3’ (reverse).

### Immunocytochemistry

Dechorionated embryos were fixed overnight at 4°C in PEM (80 mM Sodium-Pipes, 5 mM EGTA, 1 mM MgCl_2_) - 4% PFA - 0.04% TritonX100. After washing 2 × 5 min in PEMT (PEM - 0.2% TritonX100), 10 min in PEM - 50 mM NH4Cl, 2 × 5 min in PEMT and blocking in PEMT - 5% NGS, embryos were incubated 2 hrs at room temperature with primary antibodies. Following incubation, embryos were washed during 5, 10, 15 and 20 min in PEMT, blocked in PEMT - 5% NGS, and incubated again with secondary antibodies for 2 hrs. Embryos were again washed during 5, 10, 15 and 20 min in PEMT. The following primary antibodies were used: Rat@HA (Roche 11 867 423 001, 1:500). Mouse@c-Myc9E10 (Santa Cruz sc-40, 1:250). Secondary antibodies Goat@Rat-Alexa488 (Invitrogen) and Goat@Mouse-Cy5 (Jackson Immunoresearch) were used at a dilution of 1:500.

### Microscopy and image analysis

For confocal imaging, embryos were mounted in 0.75% low melting agarose (Sigma) in glass bottom dishes (Mattek). Embryos were imaged on Spinning disk (Andor) or Laser scanning confocal microscopes (Zeiss LSM710, 780 and 880) using 40x Water or 60x Oil immersion objectives. Bud stage axis extension and *in situ* gene expression patterns were documented on Leica M205FA-Fluocombi or Leica MZ-FLIII stereomicroscopes coupled to Lumenera color CCDs. Image analysis was performed using ImageJ (http://rbs.info.nih.gov/ij/). Quantifications were performed blindfolded without knowledge of the sample genotype.

### Statistical analysis

Statistical analysis was performed using R. Data normality and variance were analyzed using Shapiro-Wilk and Levene’s tests and statistical tests chosen accordingly. Complete informations about sample sizes, numerical values, and tests statistics for all experiments are provided in the Supplementary Material.

### Use of research animals

Animal experiments were performed in the iBV Zebrafish facility (authorization #B-06-088-17) in accordance with the guidelines of the ethics committee Ciepal Azur and the iBV animal welfare committee (project authorizations NCE/2013-92, 19944-2019031818528380).

## ACKNOWLEDGEMENTS

This study was supported by a CNRS/INSERM ATIP/Avenir grant, an HFSP Career Development Award (00036/2010) an ARC project grant (PJA20181208167) and the ANR DroZeMyo (ANR-17-CE13-0024-02) (MF). VMS was supported by the LABEX SIGNALIFE PhD program (ANR-11-LABX-0028-01). PS benefited from an FRM 4^th^ year PhD fellowship (FDT20140930987). AJK benefited from a 4^th^ year PhD fellowship from La Ligue Contre le Cancer. Confocal microscopy was performed with the help of the iBV PRISM imaging platform. We thank L.Bally-Cuif, I.Castanon, M.Gonzalez-Gaitan, S.Guo, C.P.Heisenberg, M.Itoh, B.Link, S.Scholpp, D.Slusarski, C.Vesque and Y.J.Jiang for the sharing of fish lines and reagents. We are grateful to R.Rebillard for excellent fish care and to T.Govekar for help with preliminary experiments.

## AUTHOR CONTRIBUTIONS

VMS, PS, AJK, SP and MF performed experiments and analyzed data. MP generated reagents used in the study. MF designed the study and wrote the manuscript.

---

**Supplementary Figure 1:**
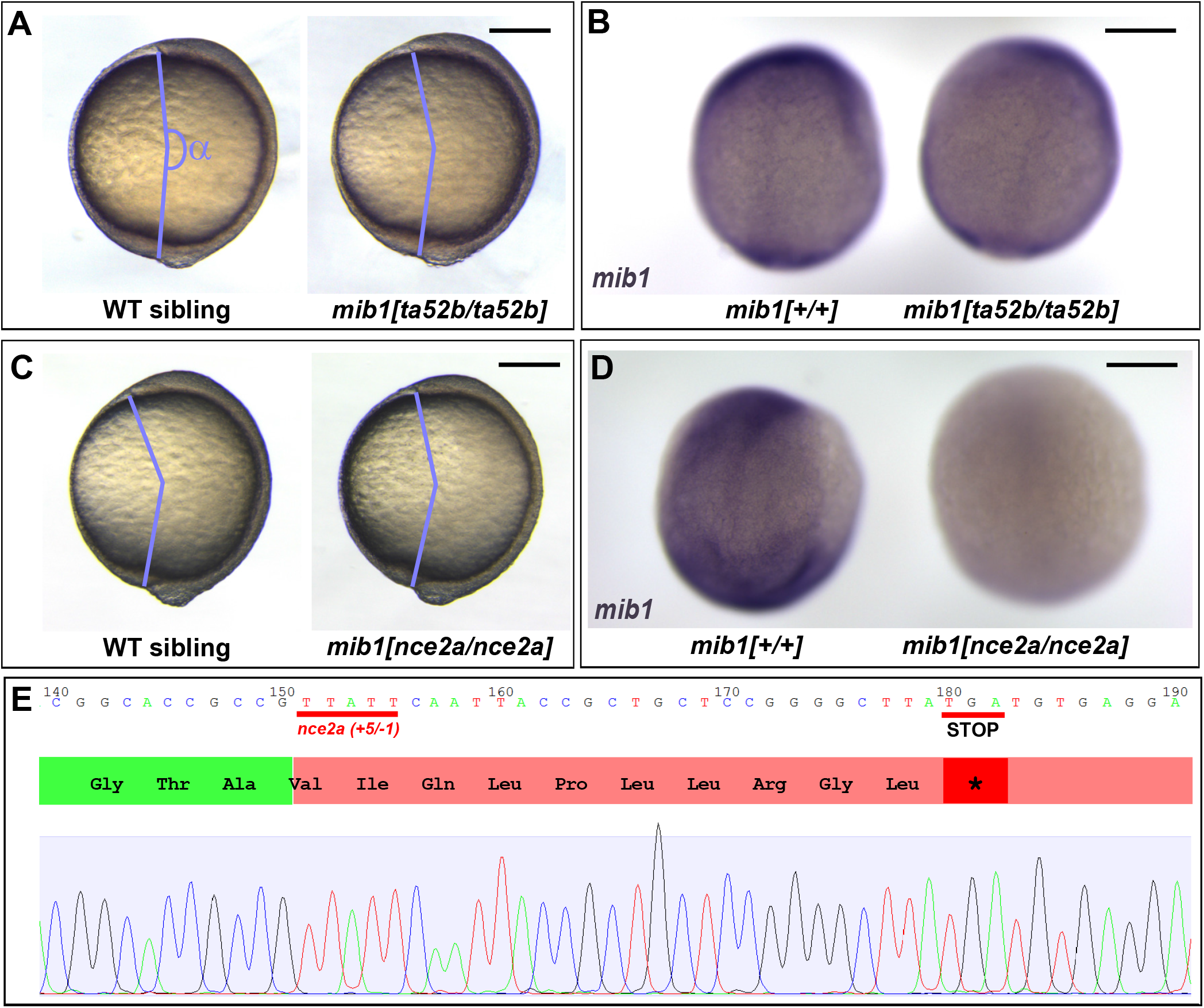
Axis extension and RNA expression in *mib1*^*ta52b*^ and *mib1*^*nce2a*^ mutants. **(A,B)** Analysis of bud stage axis extension (A) and *mib1 in situ* hybridization (B) show that the *mib1*^*ta52b*^ mutation does not impair Convergent Extension (CE) or *mib1* transcript abundance (B, n=4 mutant embryos analyzed). **(C,D)** In contrast *mib1*^*nce2a*^ mutants present a weak reduction of CE (C) and a loss of *mib1* transcripts (D, n=6 mutant embryos). **(E)** *mib1*^*nce2a*^ mutants present a CrisprCas-induced InDel that introduces a frame shift and leads to premature protein truncation. (A,C) depict lateral views of bud stage embryos, anterior up, dorsal to the right. A quantitative analysis of the corresponding data sets is provided in Fig.1D,G. (B,D) Represent dorsal views of bud stage embryos, anterior up. To warrant identical acquisition conditions, two embryos were photographed on a single picture. Scalebars: 200 µm.

**Supplementary Figure 2:**
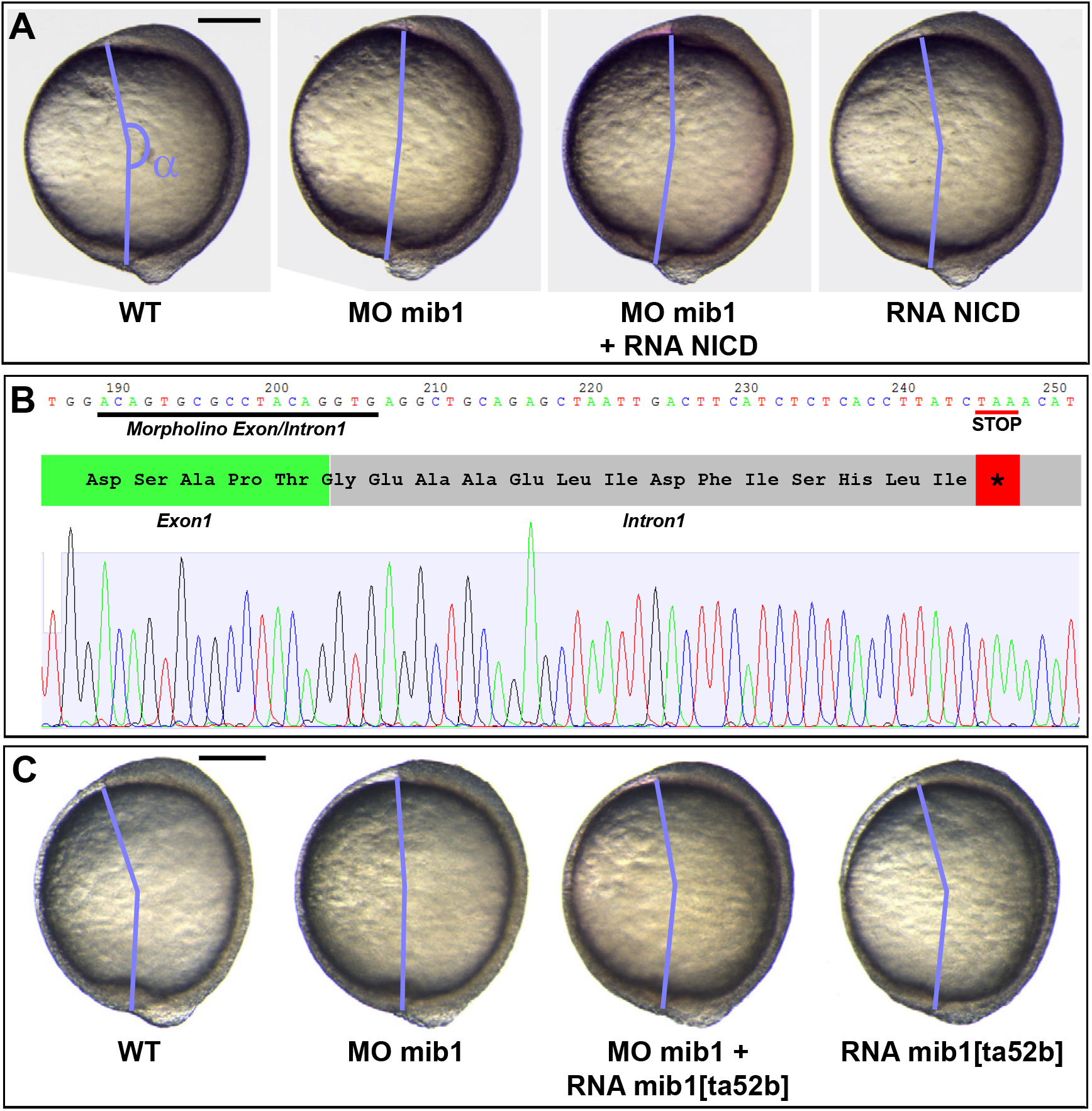
Mindbomb1 regulates convergent extension independently of Notch. **(A)** Mib1 morphants display reduced convergent extension (quantified through the measure of the axis extension level α) that is not rescued by constitutively activated Notch (NICD). **(B)** cDNA sequence from wild-type embryos injected with mib1 exon/intron1 splice morpholino. Morpholino injection causes a retention of intron 1. As a consequence, the Mib1 morphant proteins comprises only the first 76 amino acids of WT Mib1 followed by 14 intronically encoded residues and a premature Stop codon. **(C)** Mib1 morphant axis extension can be restored through the overexpression of Notch-signaling deficient mib1^ta52b^. (A,C) depict lateral views of bud stage embryos, anterior up, dorsal to the right. A quantitative analysis of the corresponding data sets is provided in Fig.1E,F. Scalebars: 200 µm.

**Supplementary Figure 3:**
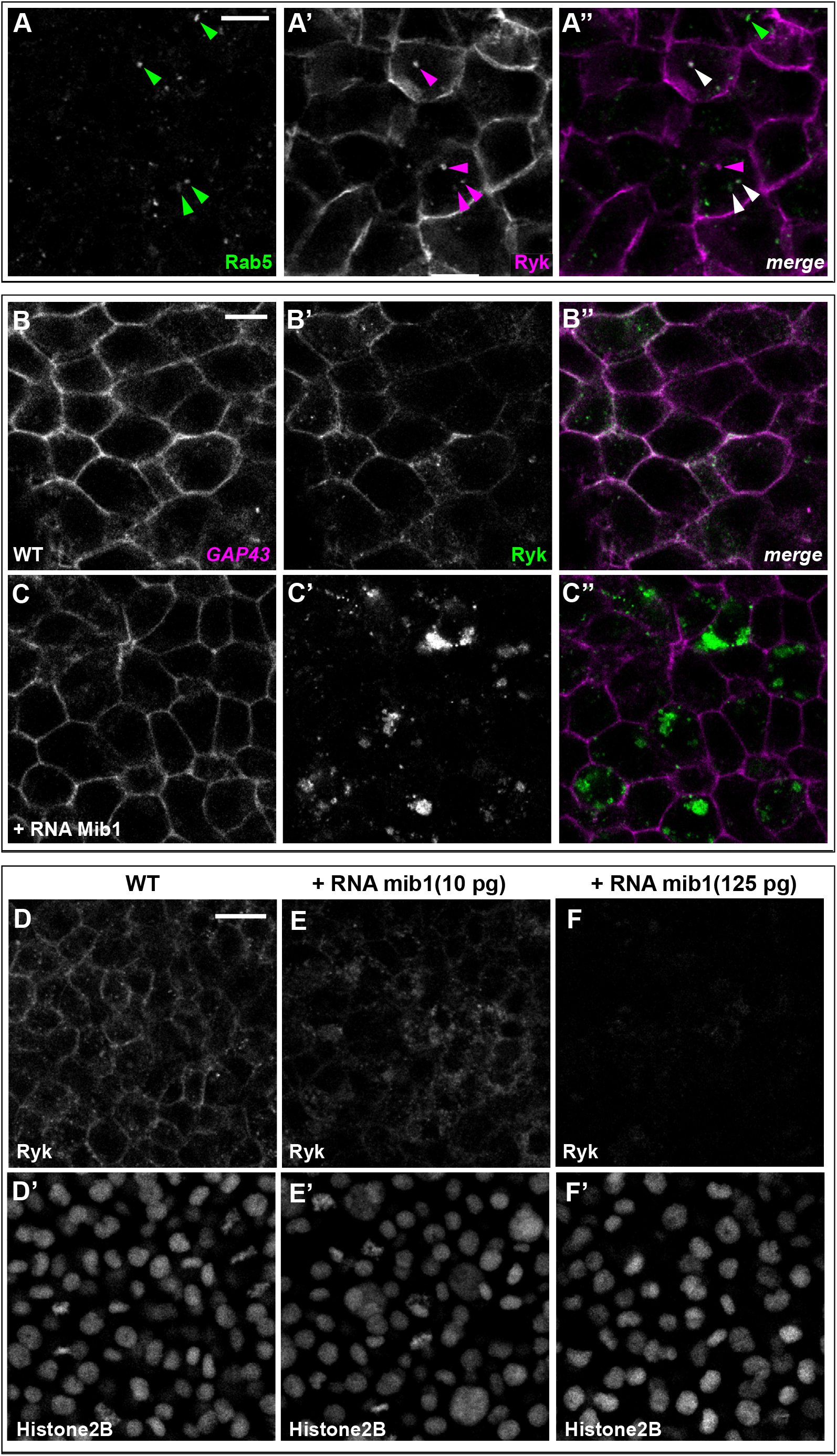
Mindbomb1 promotes Ryk internalization and degradation. **(A)** Coinjection of RNAs encoding Rab5-GFP and Flag-Ryk-Myc reveals that many Ryk-expressing intracellular compartments are positive for the early endosomal marker Rab5. Arrowheads in A’’ indicate compartments that are positive for Rab5 only (green), Ryk only (magenta), or present both markers (white). **(B,C)** Mib1 overexpression promotes the internalization of Ryk-GFP but not the one of the plasma membrane marker GAP43-RFP (n=4). **(D-F)** RNAs encoding Ryk-GFP and Histone2B-mRFP were injected with increasing amounts of mib1 RNA. While a low dose of Mib1 relocalizes Ryk from the plasma membrane to intracellular compartments (n=6), high amounts of Mib1 cause an overall loss of Ryk signal (n=6). D’-F’, The Histone2B-mRFP signal was used to ascertain that embryos had received a comparable amount of injected material. D-F and D’-F’ are sum projections of 3 consecutive slices from confocal stacks. All pictures depict dorsal views of 90% epiboly stage embryos, anterior up. Scalebars: 10 µm in A-C, 20-µm in D-F.

**Supplementary Figure 4:**
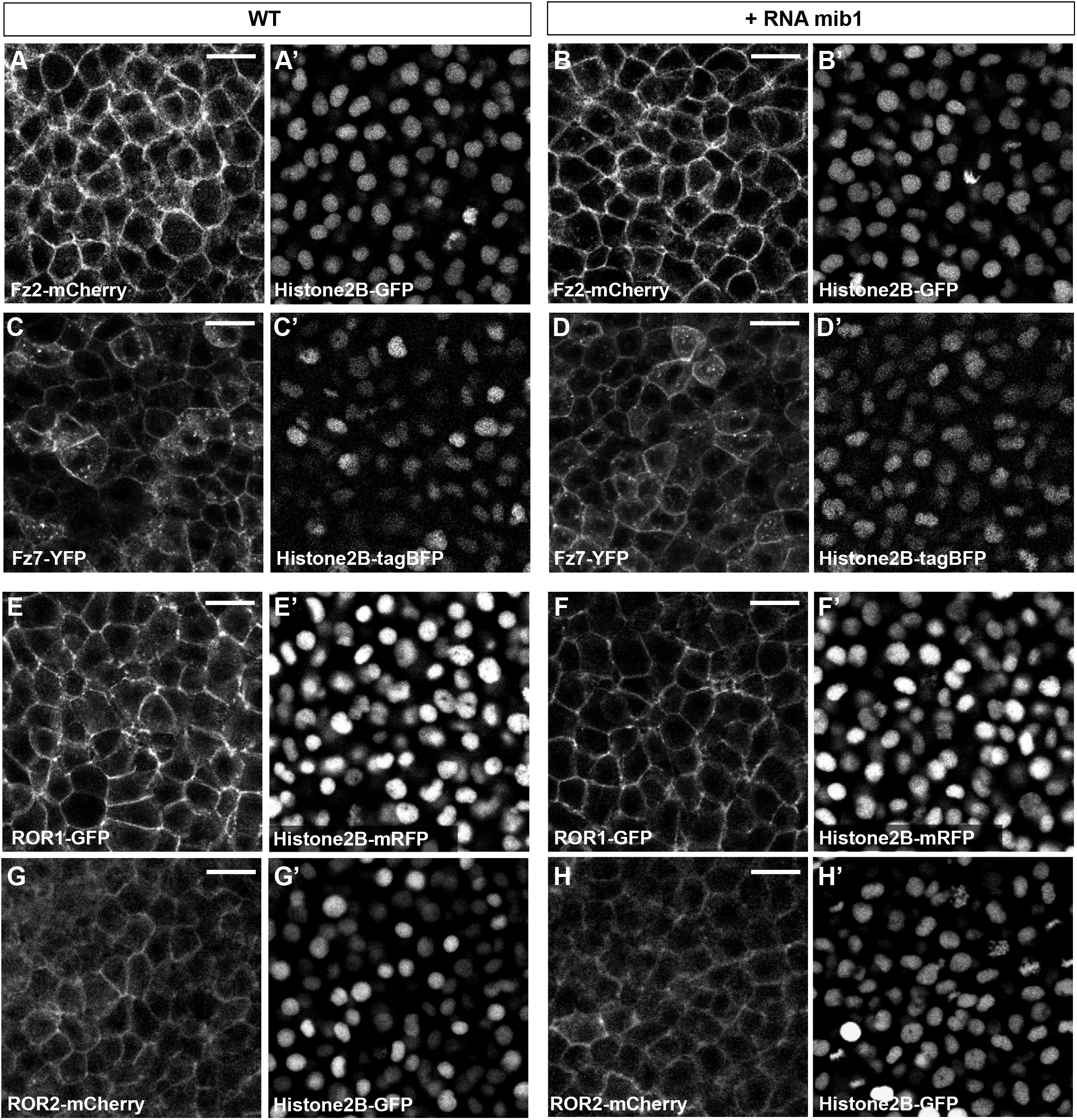
Mindbomb1 overexpression does not affect Frizzled/Ror localization. **(A-D)** Mib1 overexpression has no effect on the localization of the Wnt receptors Frizzled2 (Fz2, n=6) or Frizzled7 (Fz7, n=8). **(E-H)** Mib1 overexpression has no effect on the localization of the Wnt-binding receptor tyrosine kinases ROR1 (n=7) or ROR2 (n=10). All pictures depict dorsal views of 90% epiboly stage embryos, anterior up. G,H are sum projections of three consecutive confocal slices. A’-H’ Display the signal for fluorescently tagged Histone2B constructs that were coinjected to ascertain that control and mib1-expressing embryos had received a comparable amounts of injected material. Scalebars: 20-µm.

**Supplementary Figure 5:**
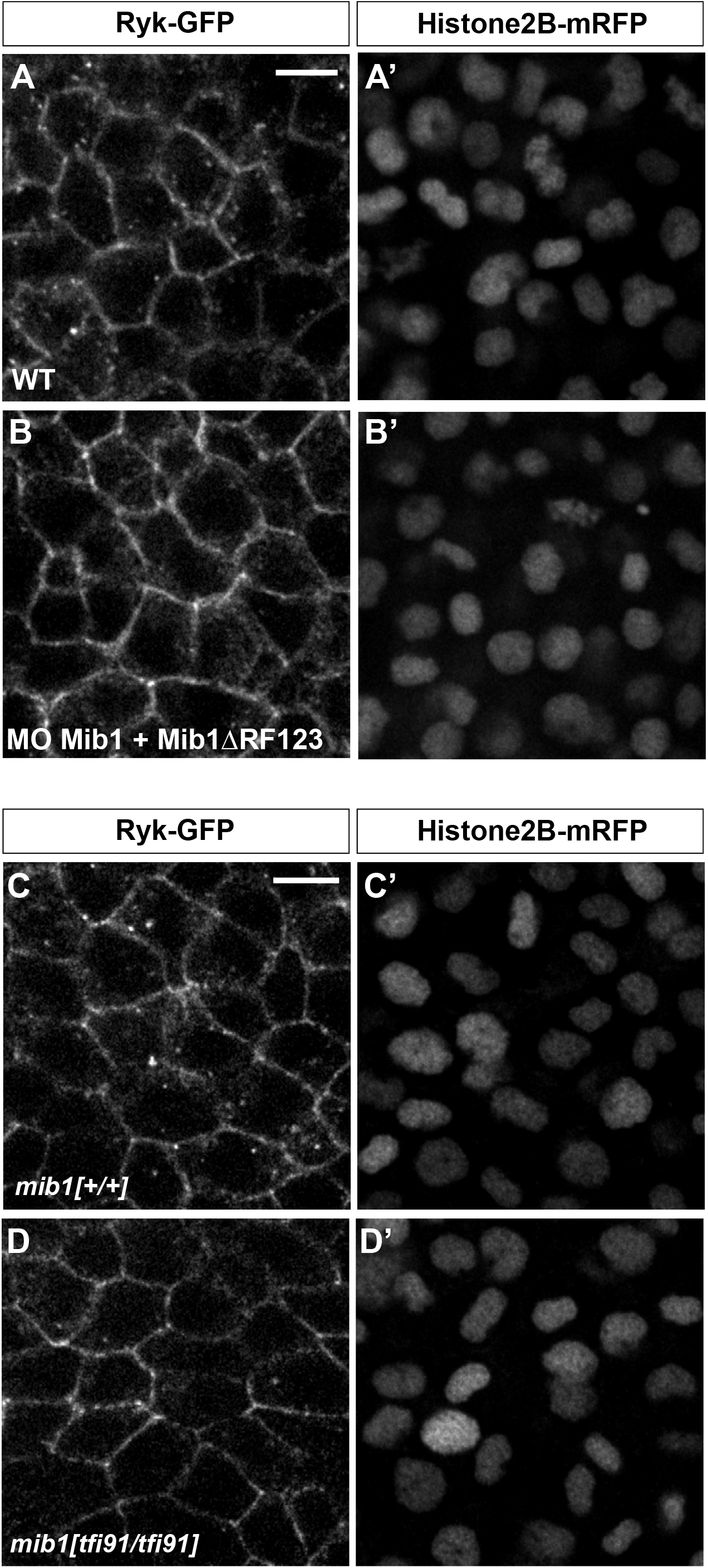
Mindbomb1 loss of function impairs Ryk endocytosis. **(A,B)** Coinjecting mib1 morpholino (MO mib1) and RNA encoding dominant-negative Mib1 (mib1ΔRF123) impairs Ryk endocytosis. **(C,D)** Ryk endocytosis is reduced in *mib1*^*tfi91*^ mutant embryos. All pictures depict dorsal views of 90% epiboly stage embryos, anterior up. A’-D’, The Histone2B-mRFP signal was used to ascertain that control and mib1-depleted embryos had received a comparable amount of injected material. Embryos depicted in A-D were injected with 12 pg Ryk-GFP RNA. A-D correspond to the display items also shown in Fig.3F-I. Scalebars: 10 µm.

**Supplementary Figure 6:**
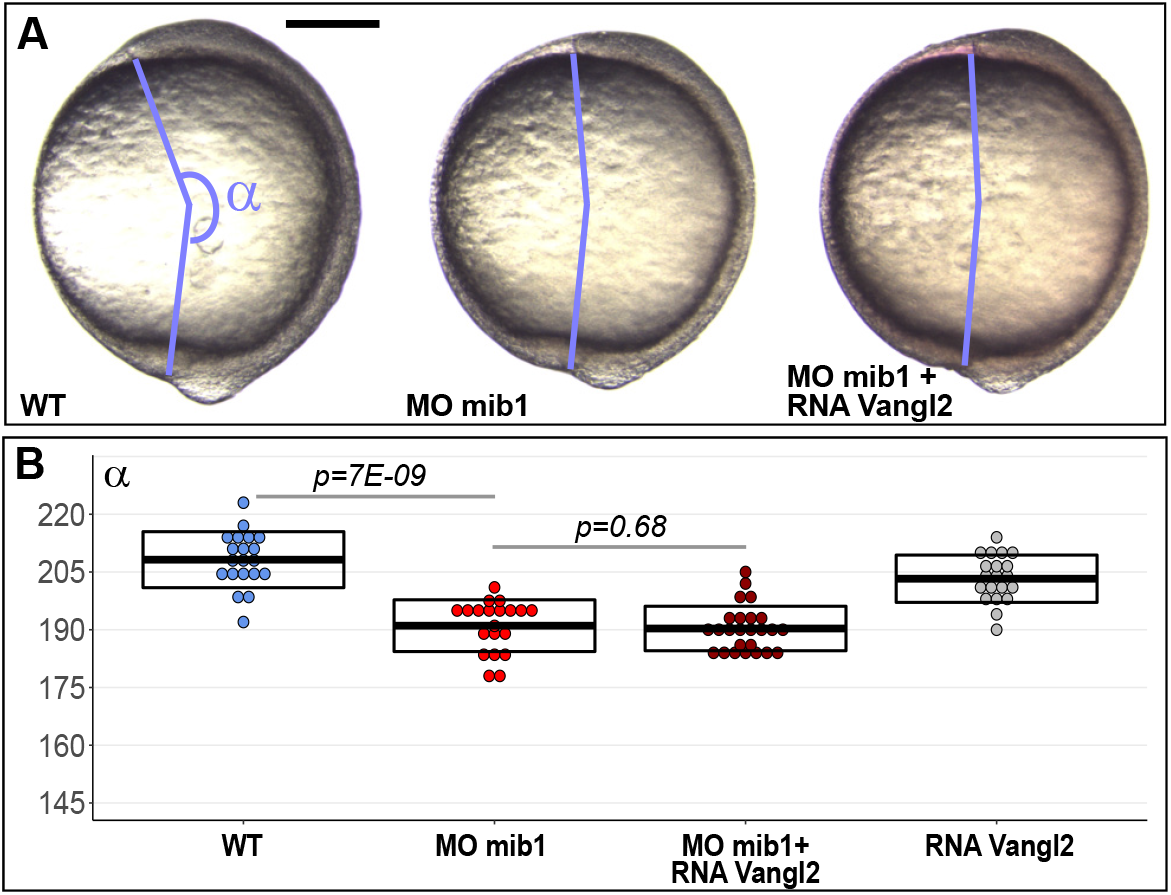
mindbomb1 morphant defects are not rescued upon vangl2 overexpression. **(A,B)** vangl2 RNA injection does not rescue axis extension in mib1 morphants. (A) Lateral views of bud stage embryos, anterior up, dorsal to the right. Scalebar 200 µm. In (B) boxes represent mean values +/- SD. See supplementary material for complete statistical information.

**Supplementary Figure 7:**
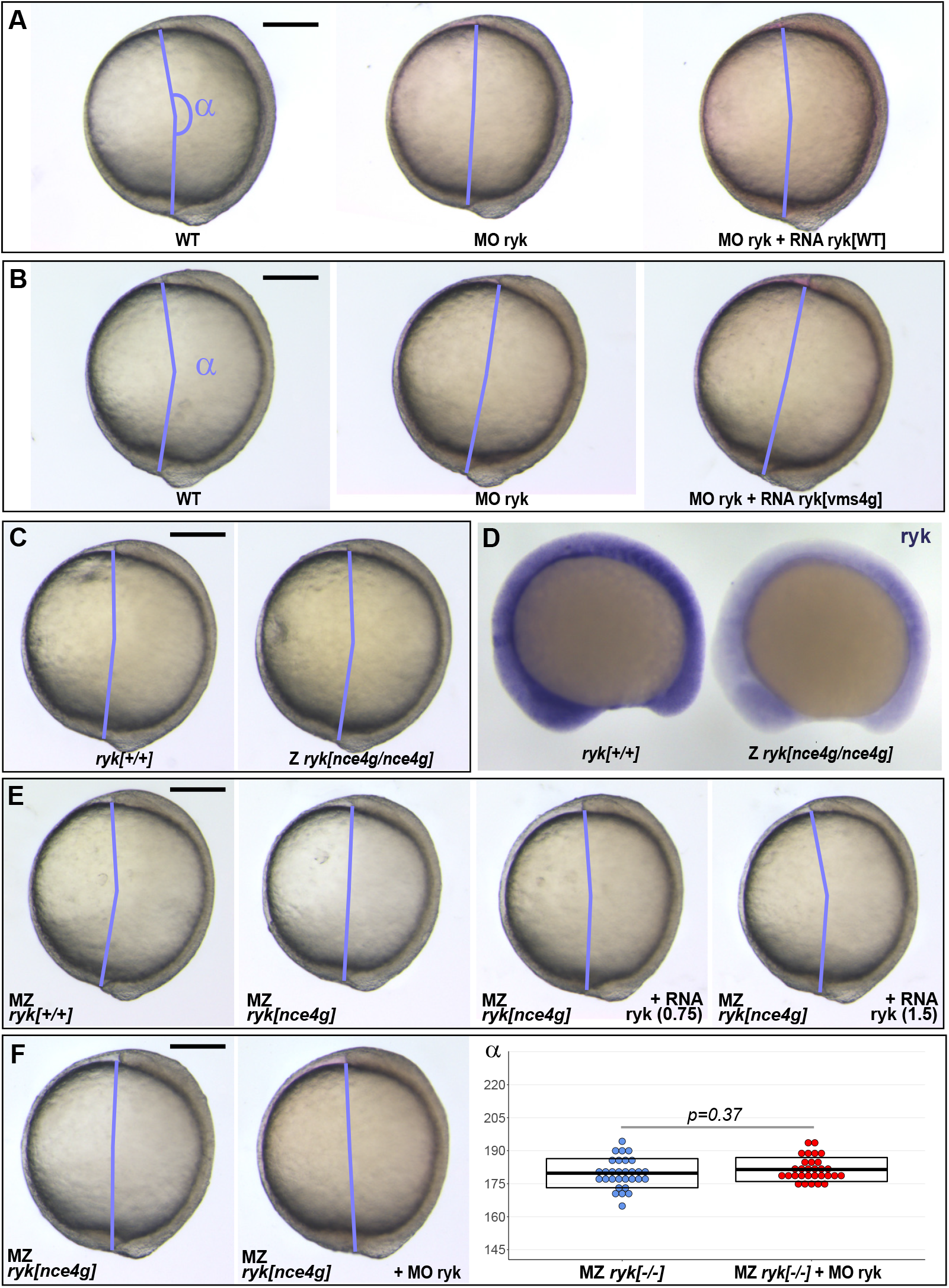
Maternal zygotic *ryk*^*nce4g*^ mutants present Convergent Extension defects. **(A,B)** ryk morphants present Convergent Extension (CE) defects that can be rescued by WT ryk (A) but not *ryk*^*nce4g*^ mutant (B) RNA. **(C)** CE is similar in Zygotic (Z) *ryk*^*nce4g*^ mutants and their WT siblings. **(D)** *In situ* hybridization reveals that *ryk* transcript levels are reduced in Z *ryk*^*nce4g*^ mutants (n=24) compared to WT siblings (n=24). 12 somite stage embryos, anterior to the left, dorsal up. To warrant identical acquisition conditions, two embryos were photographed on a single picture. **(E)** In contrast to Z *ryk*^*nce4g*^ mutants, Maternal Zygotic (MZ) *ryk*^*nce4g*^ mutants present CE defects. To exclude any defects due to genetic background variation, the parental fish used to obtain the embryos for this experiment were *ryk[+/+]* and *ryk[nce4g/nce4g]* siblings obtained from the same incross. ryk WT RNA injection allows to rescue MZ *ryk*^*nce4g*^ mutant CE defects. **(F)** ryk morpholino injection has no effect in MZ *ryk*^*nce4g*^ mutants. (A,B,C,E,F) Lateral views of bud stage embryos, anterior up, dorsal to the right. Scalebar 200 µm. In (F) boxes represent mean values +/- SD. Quantitative analysis of the data sets displayed in (A,B,C,E) is provided in Fig.4D,E,G,H respectively. See supplementary material for complete statistical information.

## SUPPLEMENTARY INFORMATION

### Complete statistical information for the experiments reported in different display items

**Fig.1A:**
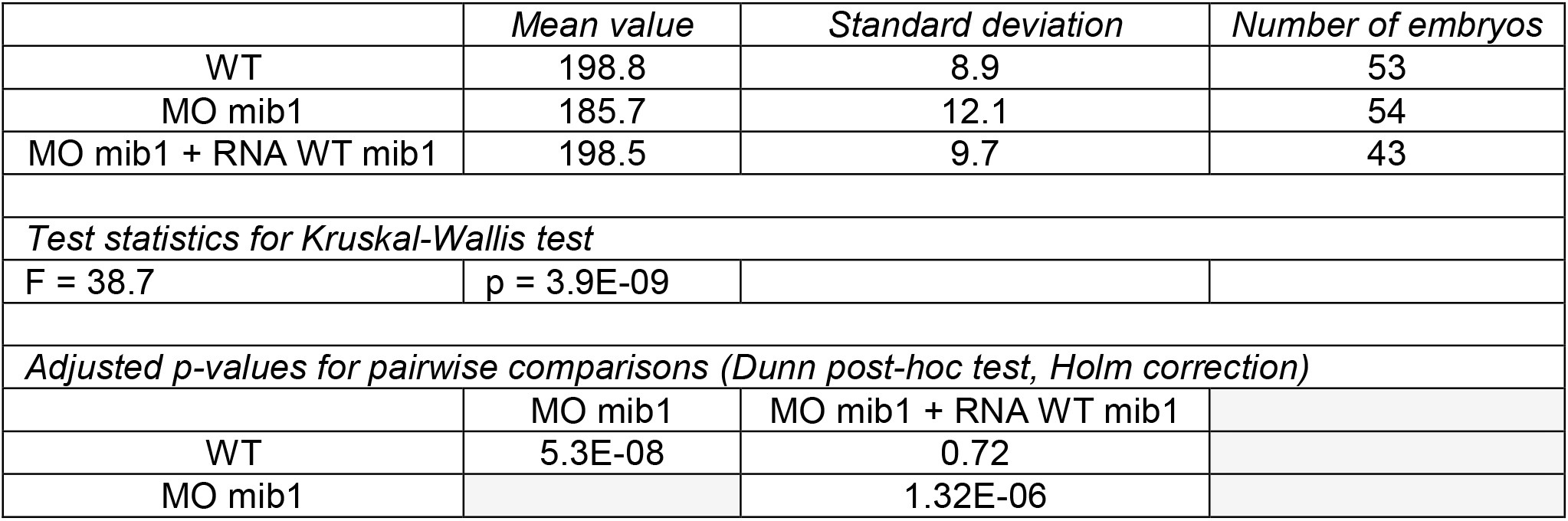
Axis extension angle in mib1 morphants.

**Fig.1B:**
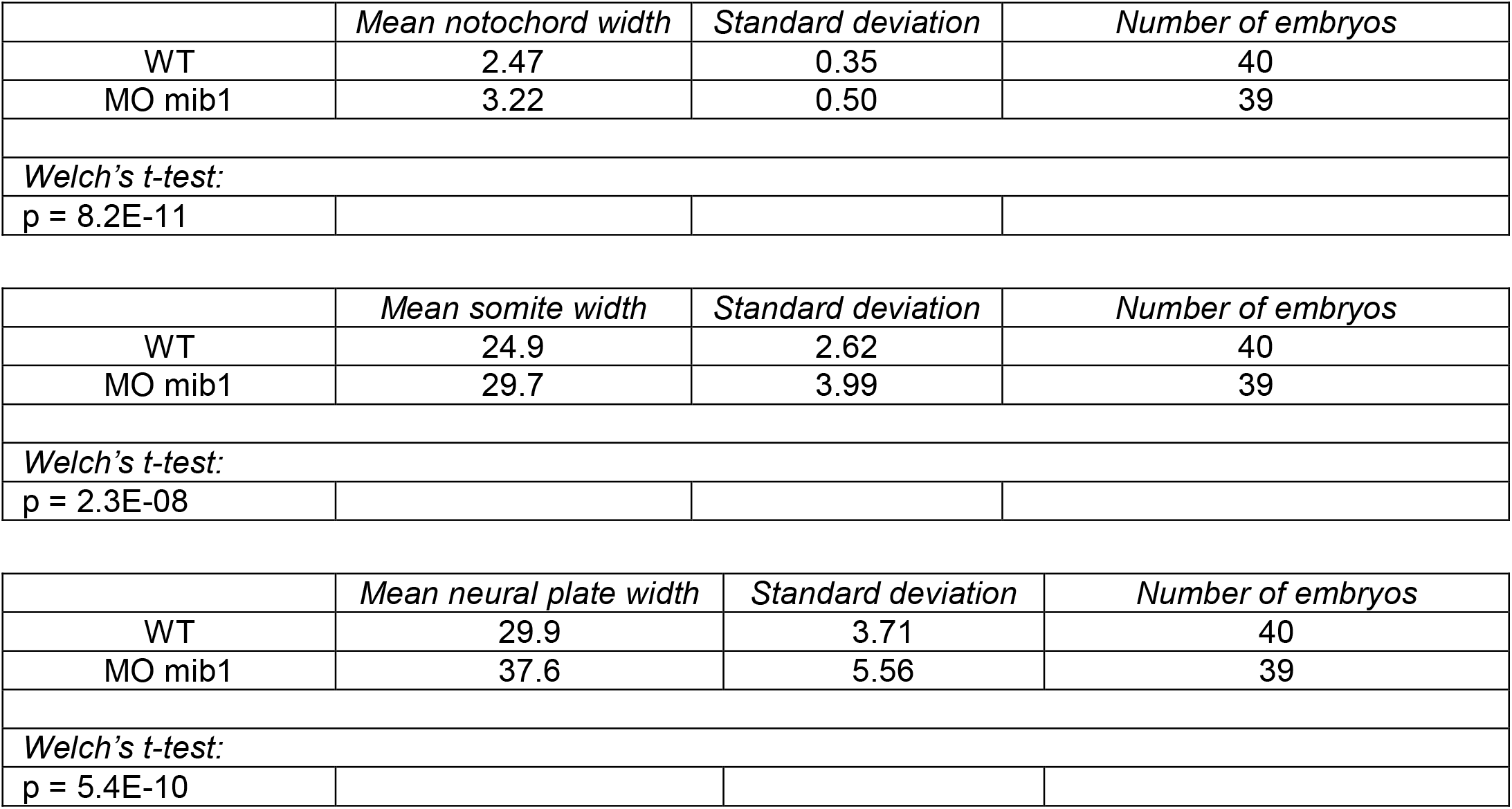
Notochord, somite & neural plate width in WT and mib1 morphants.

**Fig.1D:**
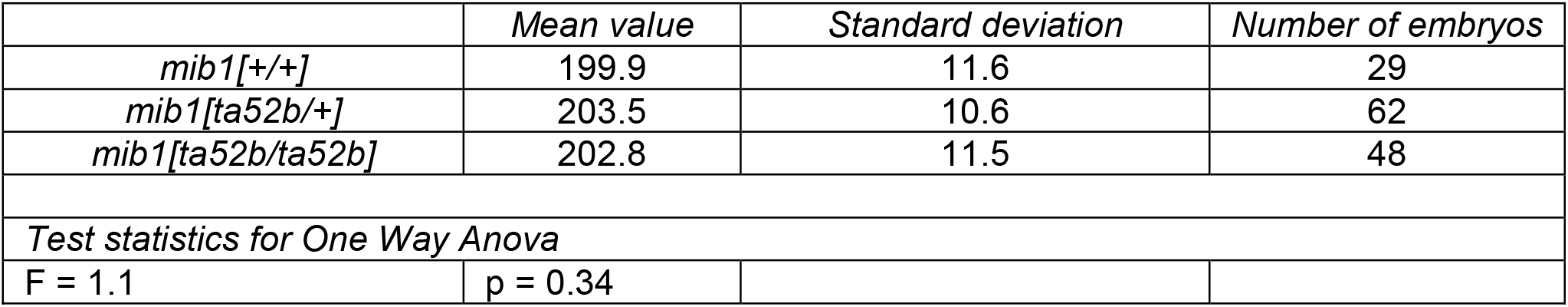
Axis extension angle in *mib1*^*ta52b*^.

**Fig.1E:**
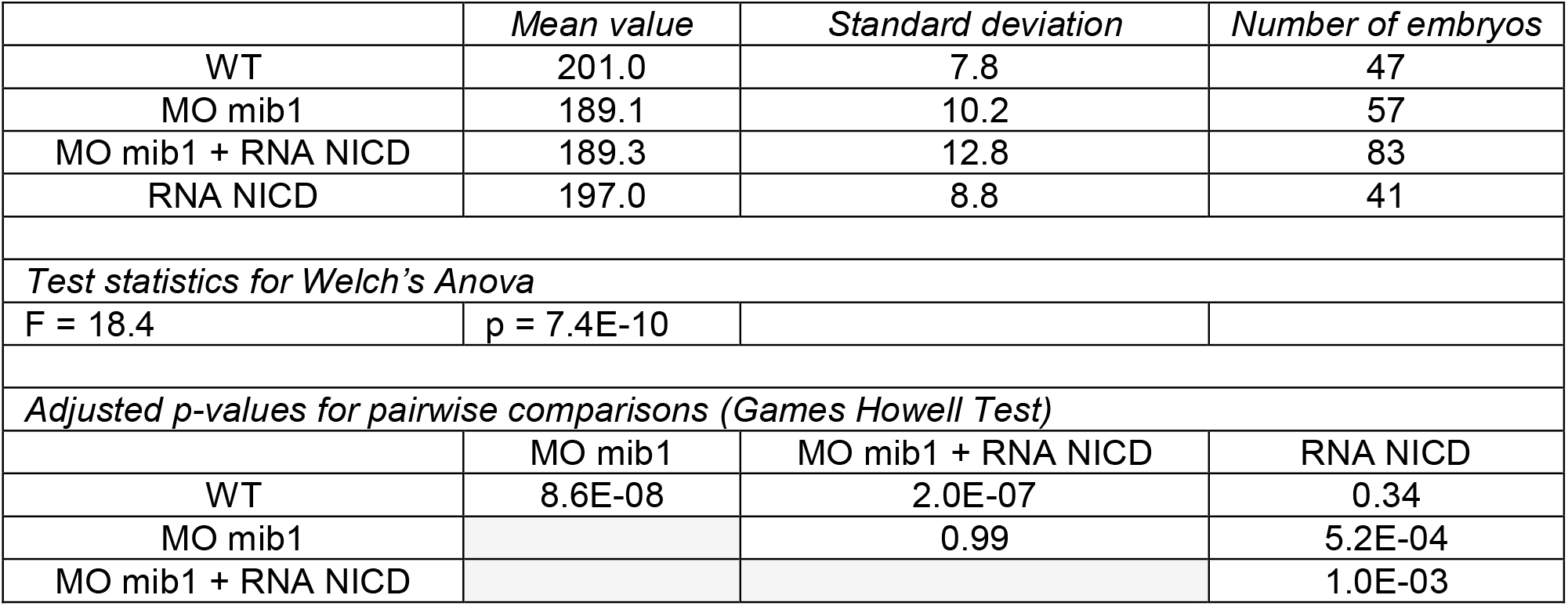
Axis extension angle in mib1 morphants injected with NICD RNA.

**Fig.1F:**
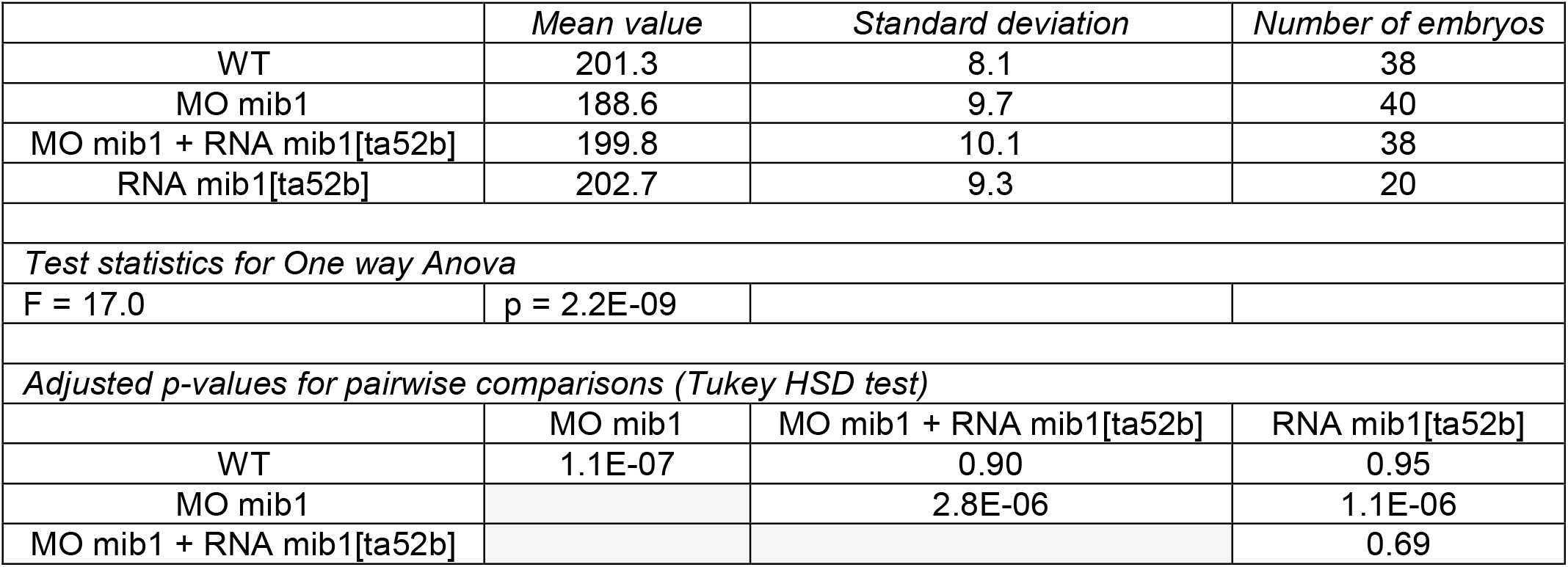
Axis extension angle in mib1 morphants injected with mib1[ta52b] RNA.

**Fig.1G:**
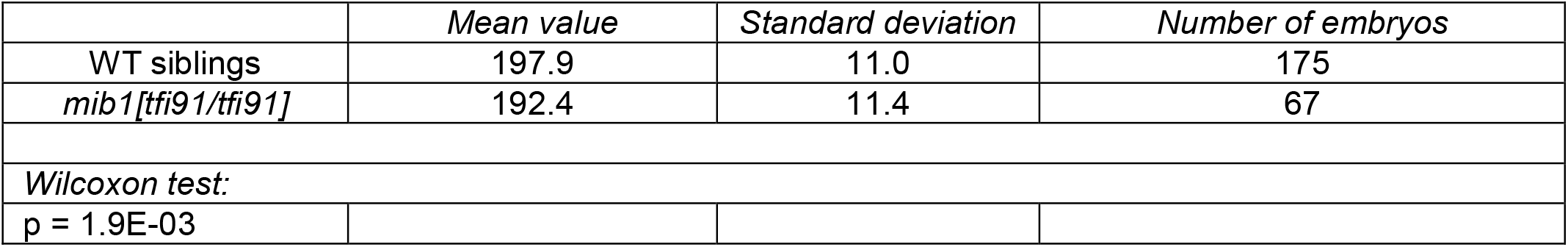
Axis extension angle in *mib1*^*tfi91*^.

**Fig.1G:**
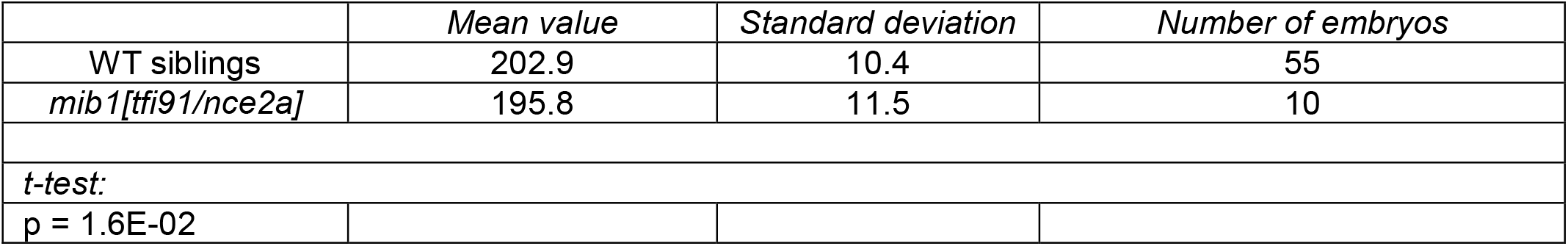
Axis extension angle in *mib1*^*tfi91/nce2a*^.

**Fig.1G:**
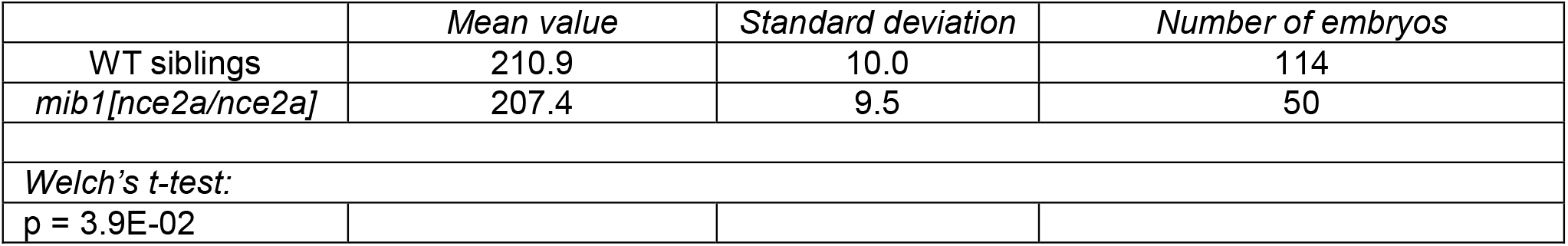
Axis extension angle in *mib1*^*nce2a*^.

**Fig.2A:**
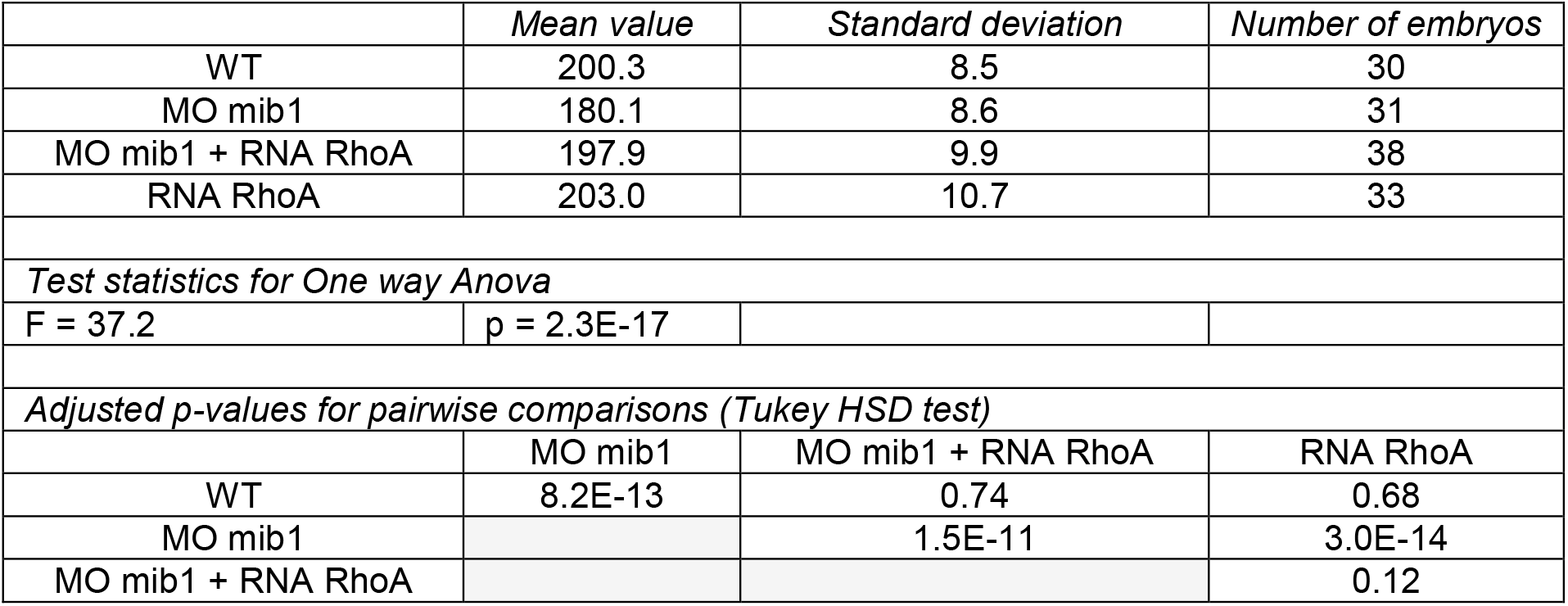
Axis extension angle in mib1 morphants injected with RhoA RNA.

**Fig.2B:**
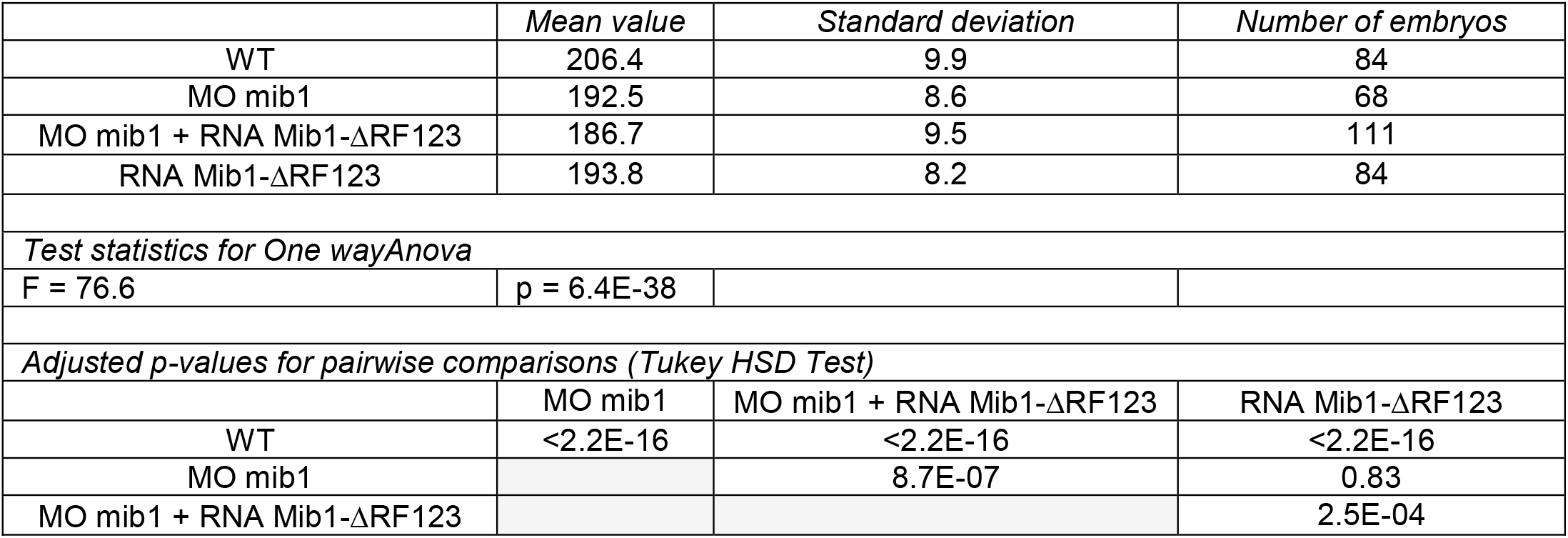
Axis extension angle in mib1 morphants injected with Mib1-ΔRF123 RNA.

**Fig.2C:**
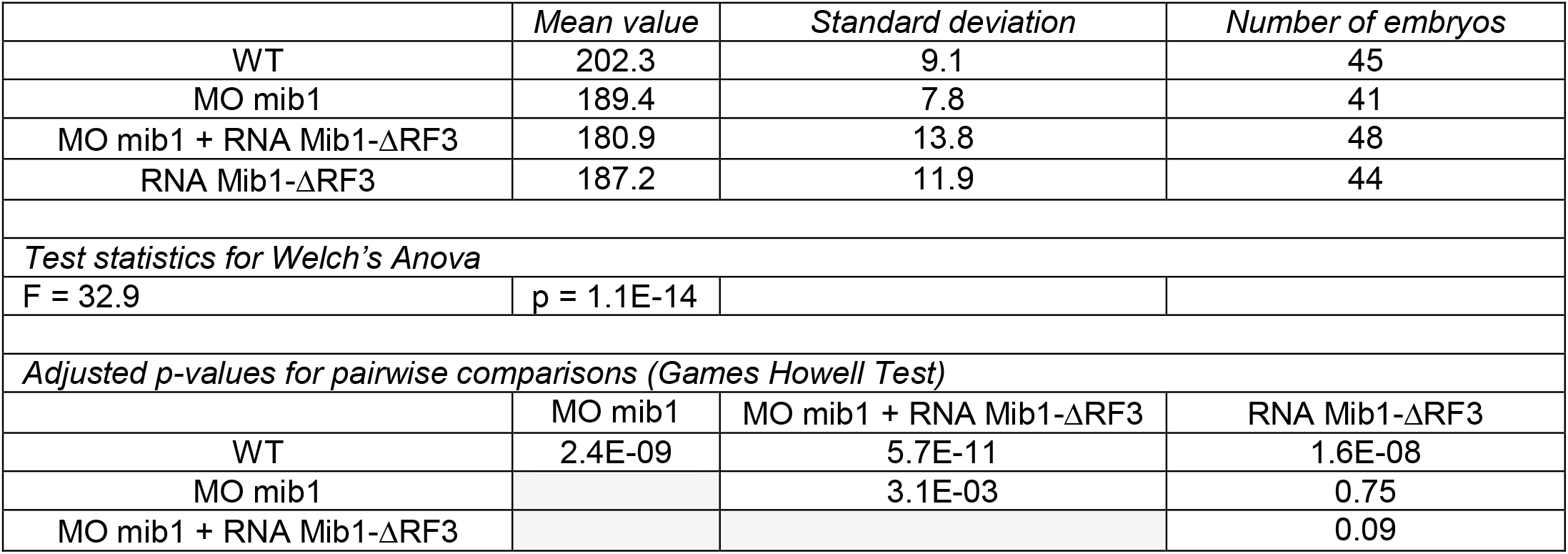
Axis extension angle in mib1 morphants injected with Mib1-ΔRF3 RNA.

**Fig.3E:**
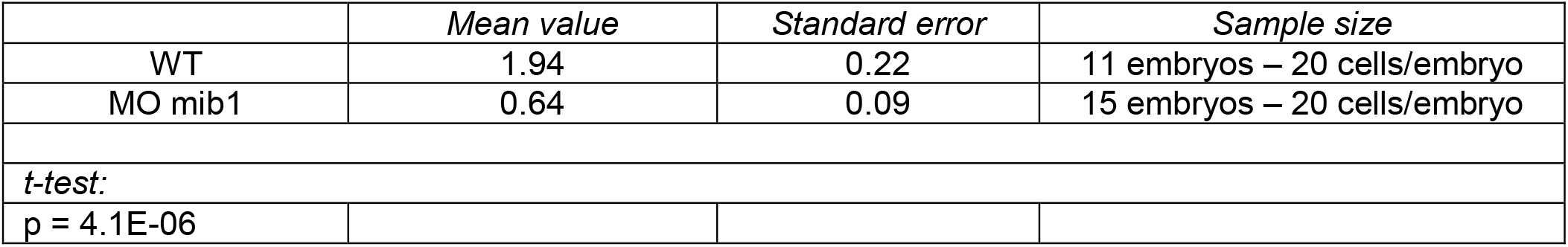
Number of Ryk endosomes in mib1 morphants injected with 3 pg Ryk-GFP RNA.

**Fig.3E:**
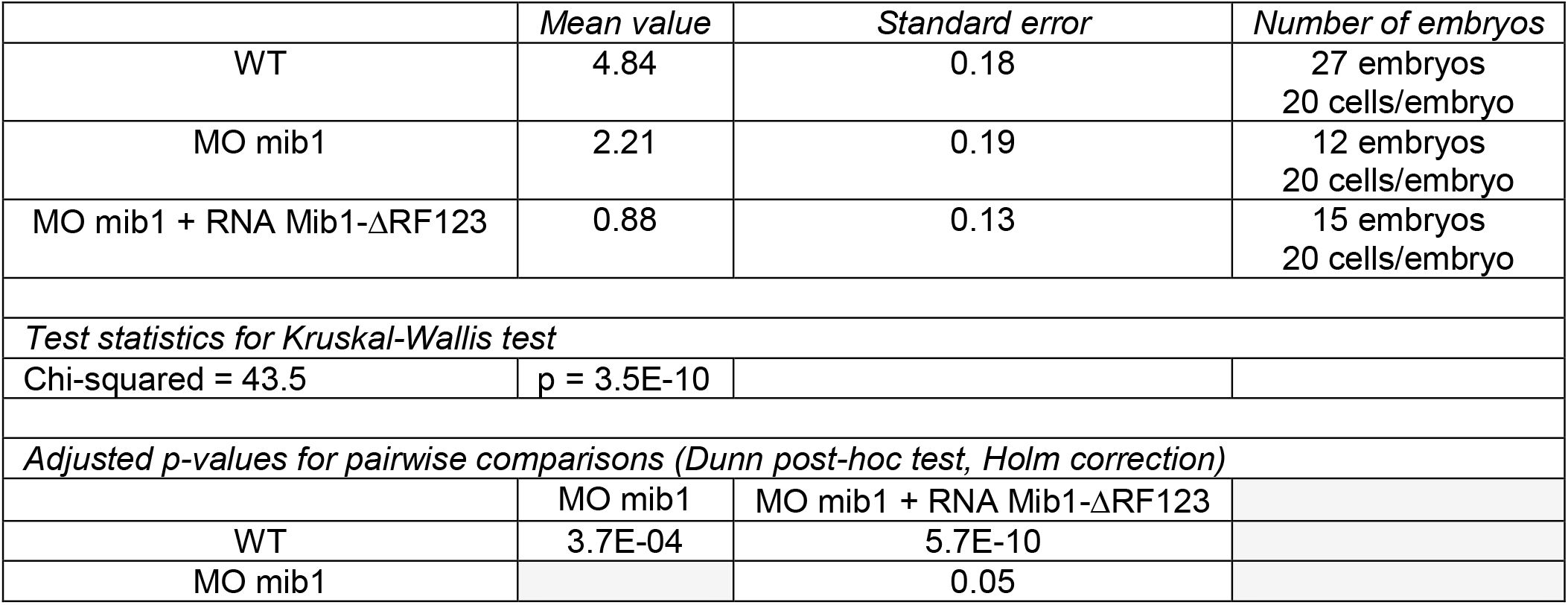
Ryk endosomes in MO mib1 + Mib1-ΔRF123 injected with 12 pg Ryk-GFP RNA.

**Fig.3J:**
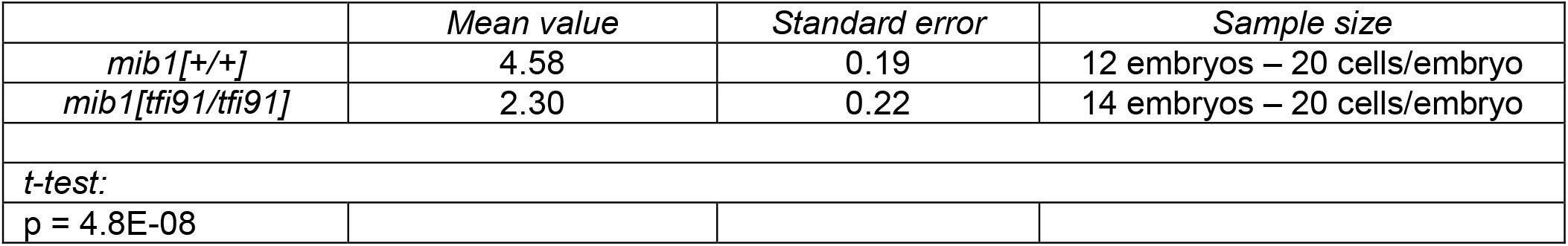
Number of Ryk endosomes in *mib1*^*tfi91*^ mutants injected with 12 pg Ryk-GFP RNA.

**Fig.3K:**
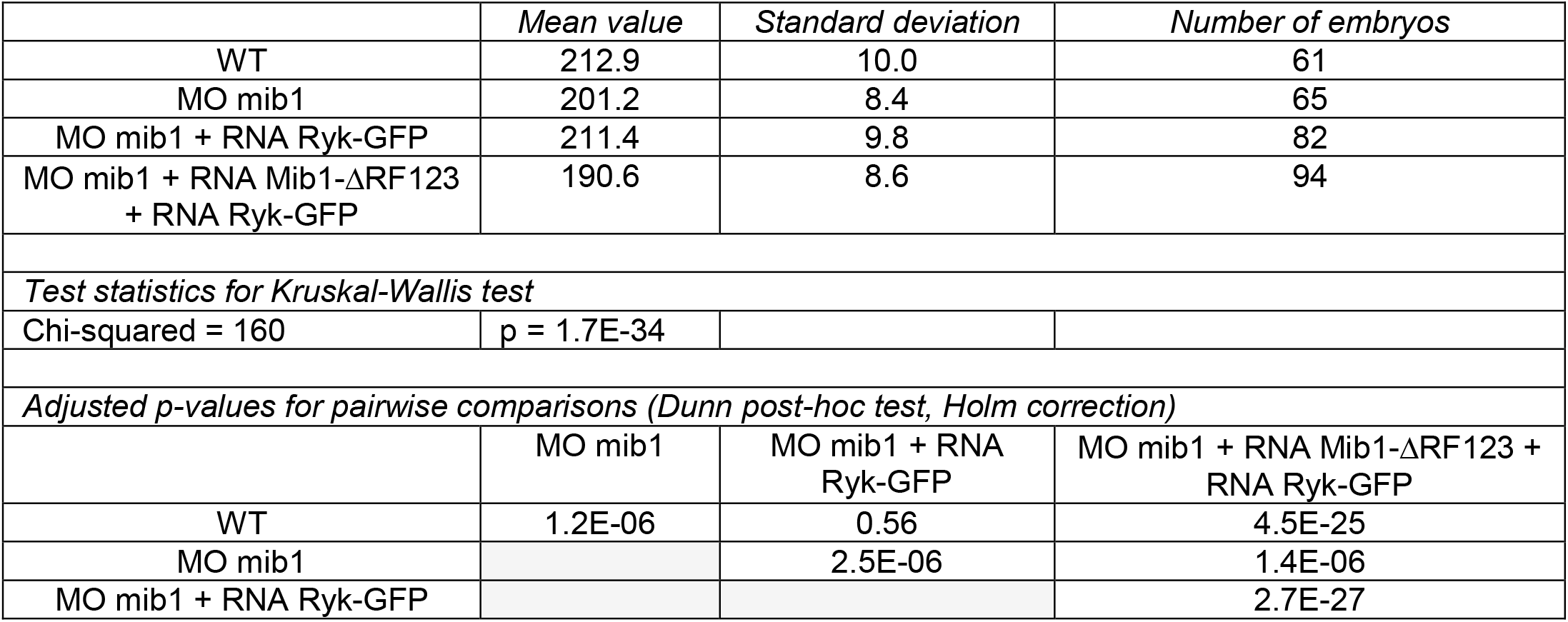
Axis extension angle in mib1 morphants injected with Ryk-GFP.

**Fig.3L:**
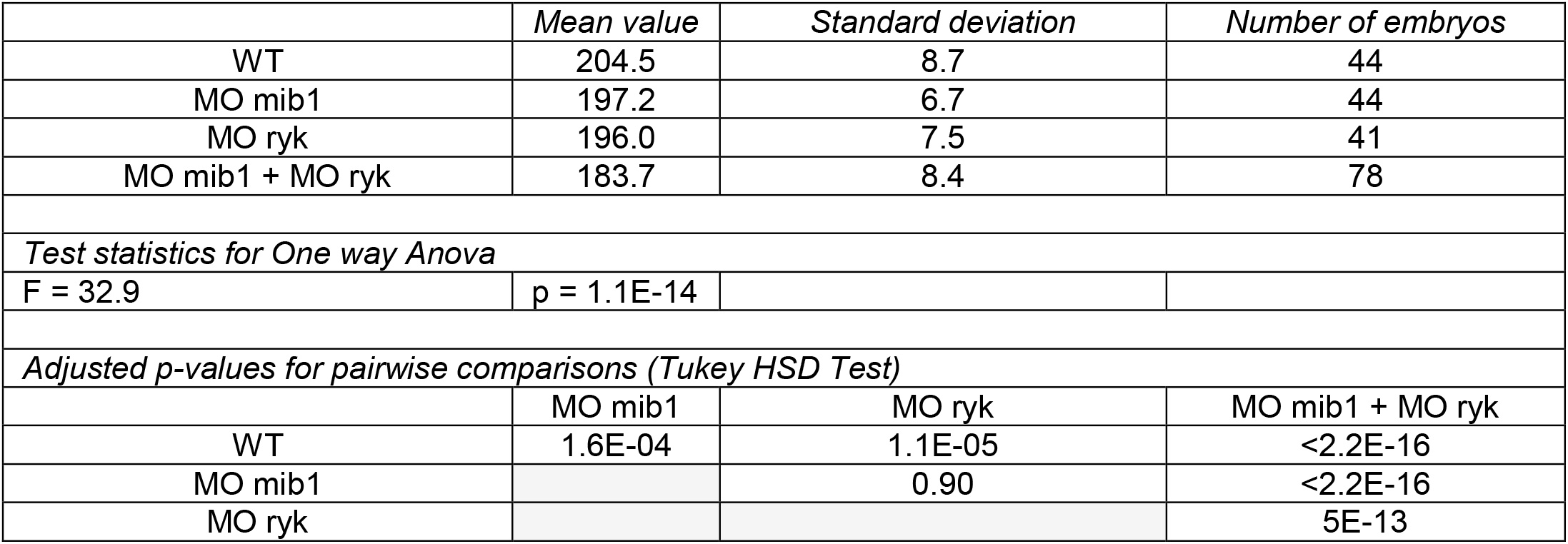
Axis extension angle in mib1 ryk double morphants.

**Fig.4D:**
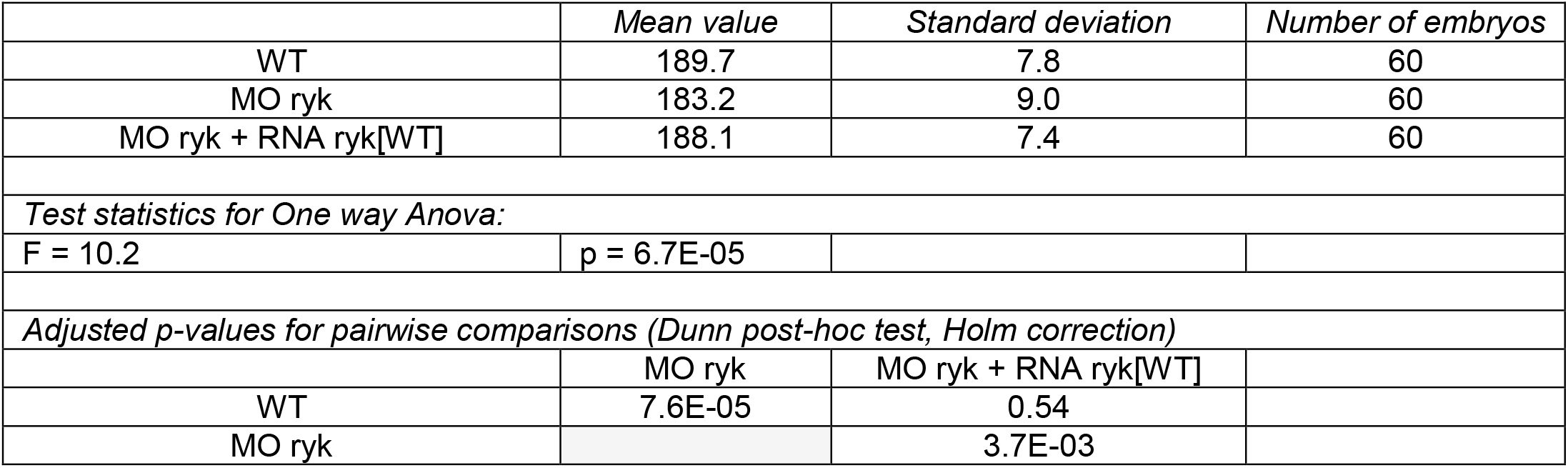
Axis extension angle in MO ryk + RNA ryk[WT].

**Fig.4E:**
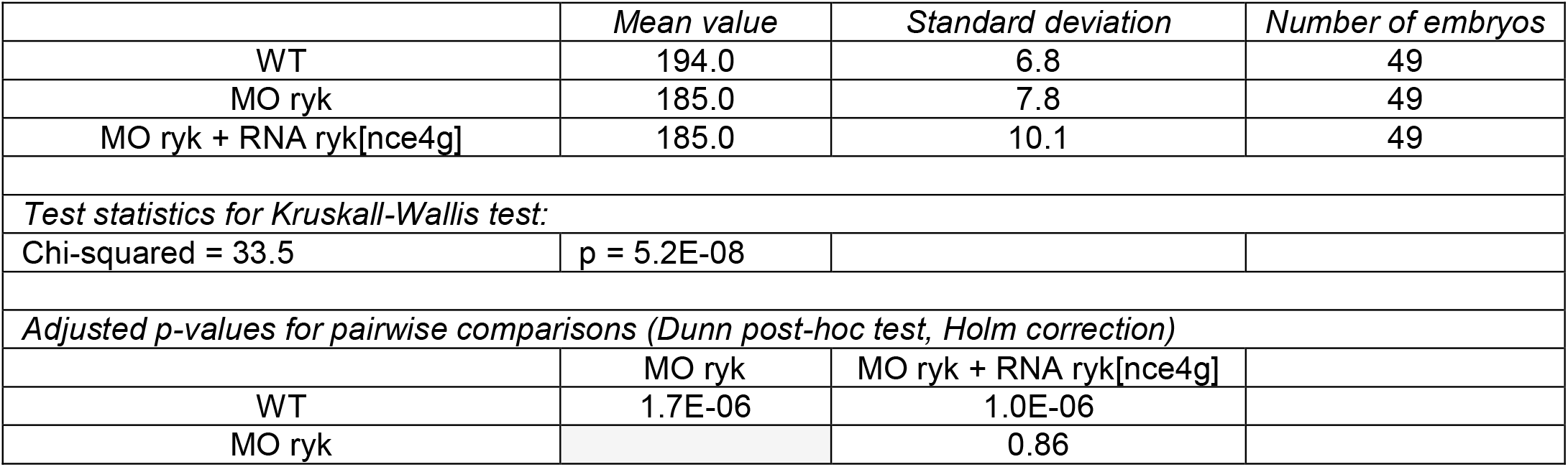
Axis extension angle in MO ryk + RNA ryk[nce4g].

**Fig.4G:**
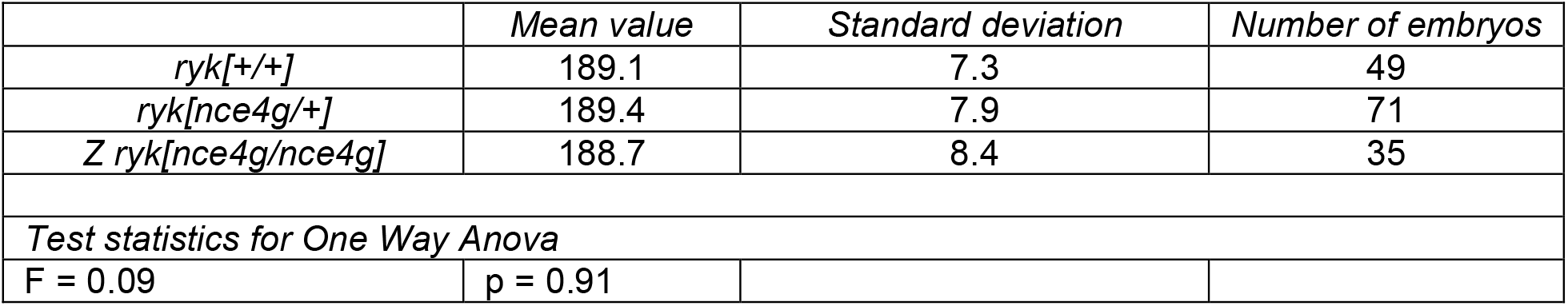
Axis extension angle in Zygotic *ryk*^*nce4g*^ mutants.

**Fig.4H:**
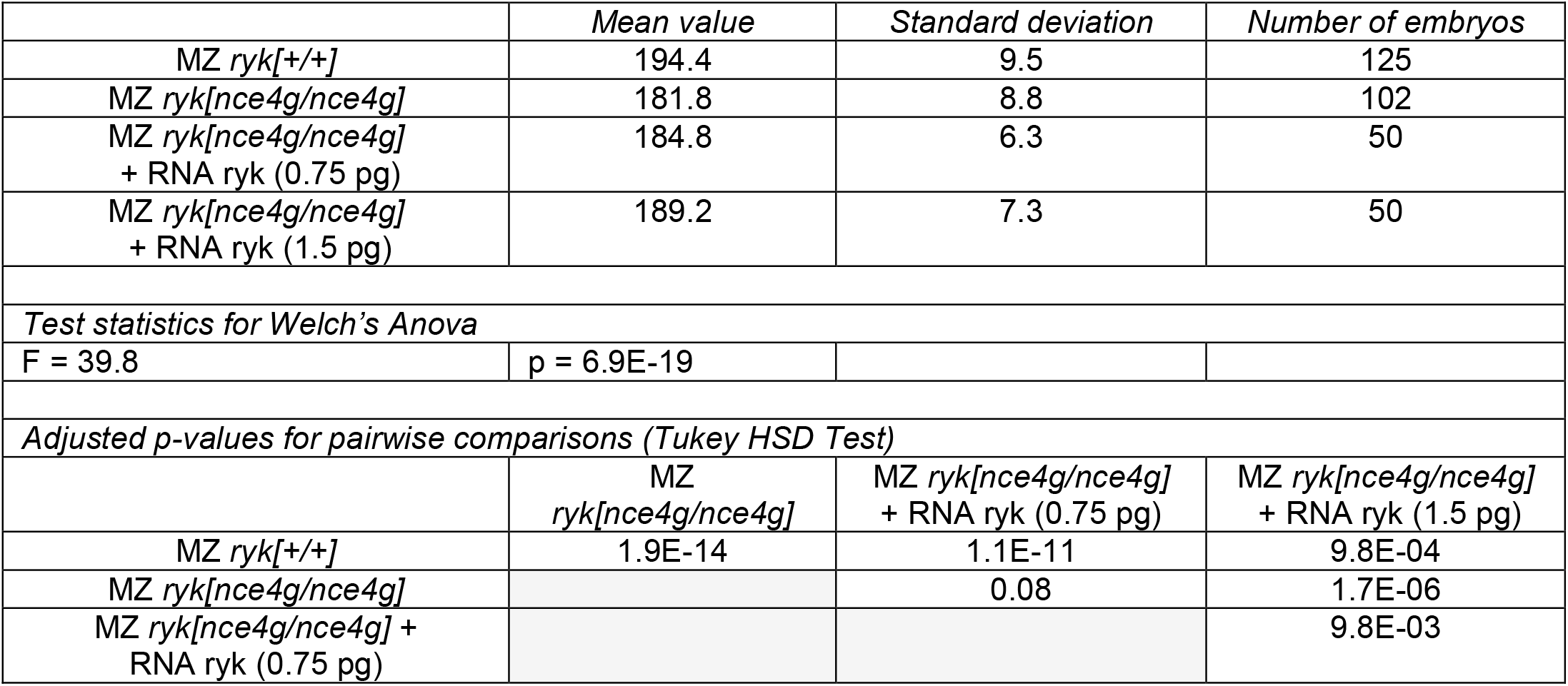
Axis extension angle in Maternal Zygotic *ryk*^*nce4g*^ + ryk RNA.

**Fig.4I:**
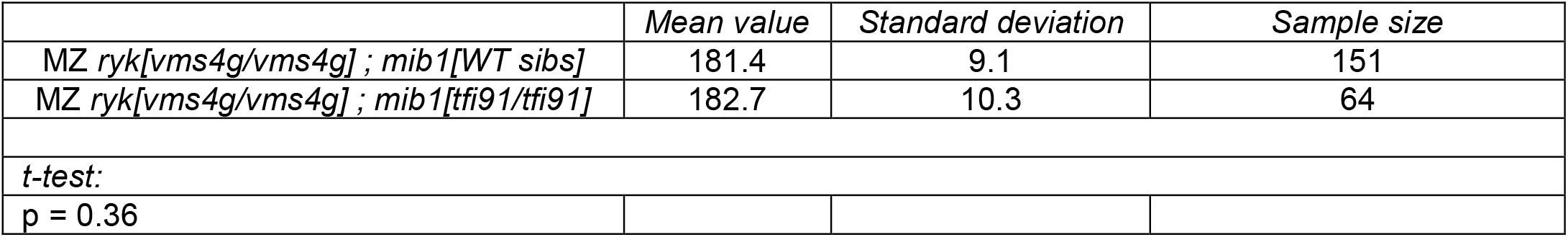
Axis extension angle in *ryk* ; *mib1* double mutants.

**Fig.S6:**
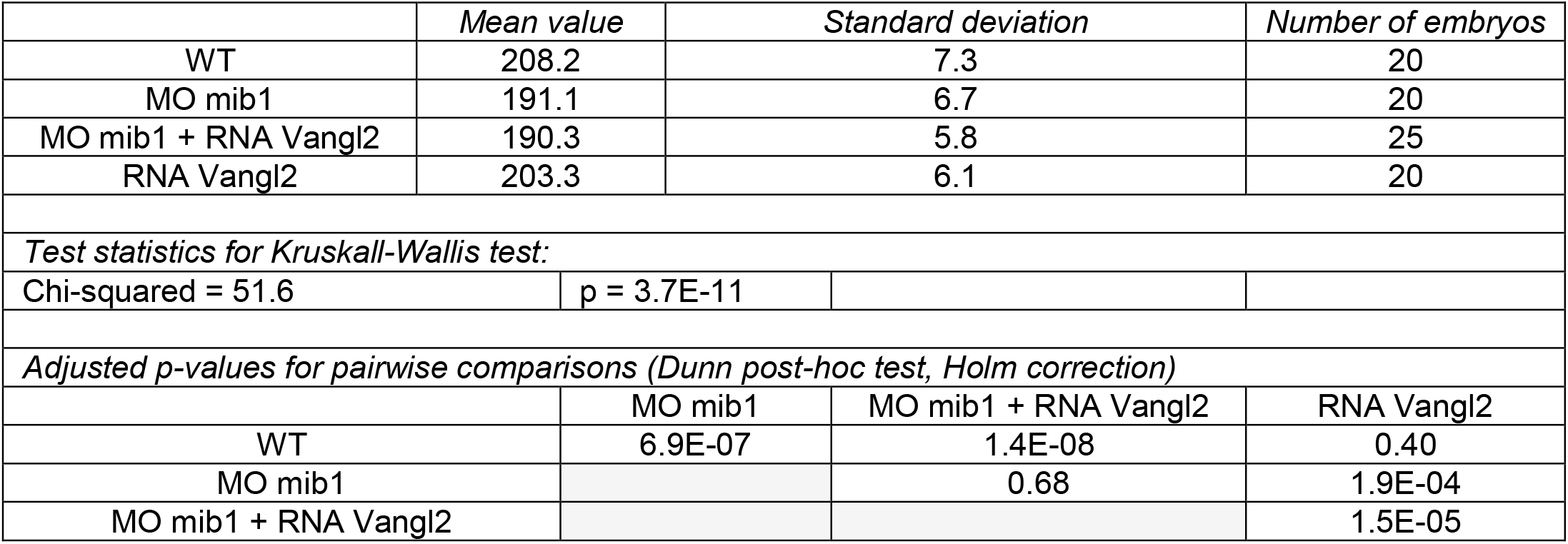
Axis extension angle in mib1 morphants injected with Vangl2 RNA.

**Fig.S7F:**
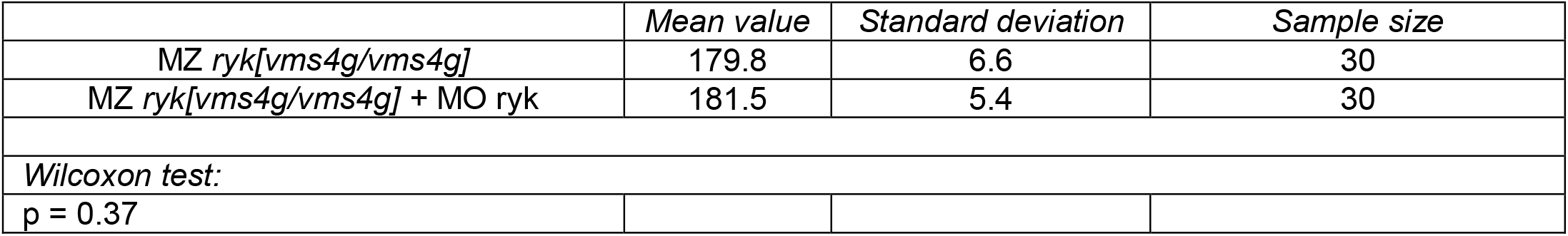
Axis extension angle in Maternal Zygotic *ryk*^*nce4g*^ + MO ryk.

